# A Synthetic Consensus Chiroptera IFN-β Induces IFN-I Signaling Across Diverse Bat Taxa

**DOI:** 10.64898/2026.10.24.761547

**Authors:** Julianne R. Bullock, Brown Bulloch, Anthony McBain, Tom Hill, Aaron Carmody, Yi He, Lucas De Sousa Paula, Paul Beare, Craig Martens, Arthur Wickenhagen, Sarah van Tol

## Abstract

Bat-associated viruses consistently spill over from bats into humans and other animals. Many bat-associated viruses, including Marburg virus and Nipah virus, cause severe disease in humans but their presentation is subclinical in bats. Advancements in tools and resources to study bat innate antiviral immune responses, including the type I IFN (IFN-I) pathway, are needed to evaluate the mechanisms that underlie differential infection outcomes between reservoir bats and spillover hosts. Commercially available IFN-Is are not bat-specific, and conspecific IFN-Is are impractical to produce for the over 1,500 described bat species. Here, we design and validate a bat order-level consensus IFN (Chiroptera IFN-β) that is functional in bat cells from six diverse bat families, including representatives from both suborders. Using Jamaican fruit bat cells, we demonstrate that Chiroptera IFN-β induces an IFN-stimulated gene repertoire comparable to conspecific IFN-β. This valuable resource can accelerate the capacity of the field to study bat innate antiviral responses and illuminate differences between bats and humans.

**Teaser:** Creation of a functional pan-bat type I interferon to enable the comparison of innate antiviral responses across diverse bat species.

## Introduction

Bats (order: Chiroptera) represent the second most speciose group of mammals, with over 1500 species [1]. Chiroptera is divided into two suborders: Yinpterochirotpera and Yangochiroptera, which diverged approximately 63 million years ago [2]. Across the two suborders, bats are divided into 21 families with diverse morphology, physiology, ecology, and behavior [3]. In addition to their vital roles in ecosystems as seed dispersers, pollinators, or predators, bats are important reservoirs of zoonotic viruses. Numerous bat species are confirmed or suspected reservoir hosts for multiple virus families (*Paramyxoviridae*, *Coronaviridae*, *Rhabdoviridae*, *Filoviridae*, and *Reoviridae*) that spill over into human populations and cause disease [4]. Spillover of bat-associated viruses, including Severe Acute Respiratory Syndrome Coronavirus 2 (SARS-CoV-2) [5], Rabies virus [6], Marburg virus (MARV) [7, 8], and Nipah virus (NiV) [9, 10], continues to threaten human and animal health in the Anthropocene.

Many of these bat-borne viruses show high virulence in spillover hosts despite lacking apparent immunopathology or mortality in bats [11–19]. The mechanisms underlying the differential pathogenicity between bats and spillover hosts are not well defined. Differences in viral activation, or lack thereof, of the innate antiviral immune response among species may be key in determining whether clinical disease develops in response to infection.

Activation of the type I interferon (IFN-I) response and signaling pathway is critical to regulate early responses to virus infection. Host detection of viral products, such as double-stranded RNA by pattern recognition receptors (PRRs) retinoic acid-inducible gene I (RIG-I) and Melanoma Differentiation-Associated gene 5 (MDA5), triggers signaling cascades that activate the transcription and secretion of IFN-Is and other cytokines [20, 21]. Secreted IFN-Is bind to the IFN-I receptor (IFNAR) and activate the nuclear translocation of transcription factors, Signal Transducer and Activator of Transcription 1 (STAT1) and 2 (STAT2), to induce expression of interferon-stimulated genes (ISGs) which establish an antiviral microenvironment [22]. The IFN-I pathways are conserved across vertebrates, and many viruses have evolved strategies to block IFN-I induction and signaling to enable completion of their replication cycle.

Despite the conservation of the major components of the IFN-I signaling pathways among vertebrates, IFN-I and IFNAR amino acid sequences are highly diverse. This diversity in the IFN-I and IFNAR amino acid sequences results in species-specific interactions [23]. Due to this constraint, the field has either generated conspecific IFN-Is or used a commercially available universal IFN-I (U-IFN) to evaluate the IFN-I signaling pathway in new species. U-IFN is a synthetic human IFN-α hybrid protein that induces IFN-I signaling across diverse mammalian cells [24], including some bat species [25–27]. A potential concern with U-IFN is that it may not induce an authentic ISG repertoire compared to conspecific IFN-Is. Since Chiroptera includes over 1,500 species, generating and validating conspecific, recombinant IFN-I for each species is infeasible. Engineering an order-wide consensus IFN-I is a potential approach to bypass the need to synthesize conspecific IFN-βs for each species of interest.

Here, we design and validate Chiroptera (Chi) consensus IFN-β as a proof-of-concept. We demonstrate that Chi IFN-β triggers IFN-I signaling that is antiviral and sensitive to IFN-I signaling inhibitors in cell lines representing six bat family’s across both suborders. Overall, these results support that Chi IFN-β stimulates an authentic IFN-I signaling response that can be applied to study the innate antiviral response of diverse bat taxa.

## Results

### Generation of conspecific, Phyllostomid consensus, and Chiroptera consensus IFN-βs

To assess the capacity for an order consensus IFN-β to stimulate IFN-I signaling across diverse bat species in comparison to conspecific IFN-β, we first designed a Chiroptera (Chi) consensus IFN-β based on available bat *Ifnb1* gene sequences. At the time of the design of Chi IFN-β, sequences from 46 bat species across ten bat families, including representatives from both the Yangochiroptera (6) and Yinpterochiroptera (4) suborders, were available (Supplementary Table 1, Supplementary Fig. 1A). Family-level consensus *Ifnb1* sequences were assembled first, then a Chi *Ifnb1* consensus sequence was generated from the ten family-level consensus sequences. This approach prevented bias toward bat families with higher sequence availability, such as Phyllostomidae (Phy), Vespertilionidae, and Pteropodidae. When only two species were present in a bat family, the consensus sequence included degenerate nucleotides.

To enable comparison of Chi IFN-β function, *Artibeus jamaicensis* (Art) *Ifnb1*, the Phy consensus *Ifnb1*, and Chi *Ifnb1* were synthesized. An interleukin-2 (IL-2) secretion signal, His-tag, and Tobacco Etch Virus (TEV) protease cleavage site were added upstream of each *Ifnb1* open reading frame to enable secretion and cleavage of the His-tag following expression of the mature form of IFN-β (Fig. 1A). Following expression, purification, and removal of the His-tag (Supplementary Fig. 1B), the IFN-β monomer-containing fractions were collected and analyzed with SDS-PAGE (Supplementary Fig. 1C).

**Fig 1.**
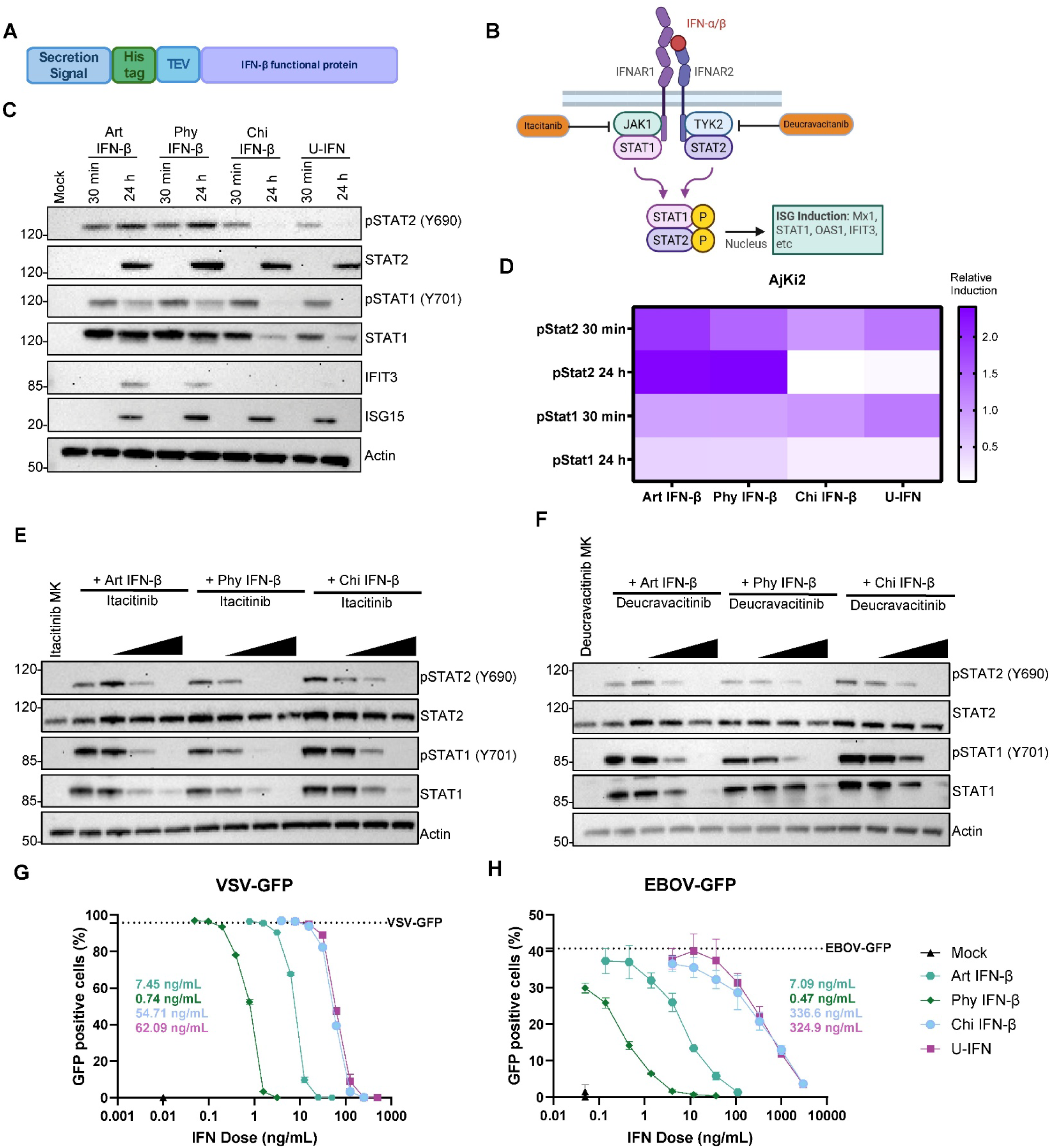
Conspecific, Phyllostimid consensus, and Chiroptera consensus IFN-β induce type-I IFN signaling in Jamaican fruit bat cells. (A) Schematic of the IFN-β recombinant plasmid used to generated *Artibeus jamaicensis* (Art), Phyllostomid consensus (Phy), and Chiroptera consensus (Chi) interferon-beta (IFN-β) including the secretion signal, a His-tag for purification, and a Tobacco Etch Virus (TEV) protease cleavage site. (B) Overview of the JAK-STAT pathway utilized by type-I IFNs to induce ISGs through intermediate STAT signalers. Itacitinib and Deucravacitinib inhibit type-I signaling through interactions with JAK1 and TYK2, respectively. (C) Immunoblot for AjKi2 cells untreated (mock) or treated with 500 ng/mL of Art IFN-β, Phy IFN-β, and Chi IFN-β or U-IFN and lysed after 30 minutes (min) or 24 hours (h). Shown western blot representative of three independent experiments. (D) Densitometric quantification of immunoblot bands normalized to actin and Chi IFN-β at 30 min to show relative induction. Quantification based on densitometry from three independent experiments. (E, F) Immunoblots for AjKi2 cell treated for 24 hours with three concentrations (1, 5, 25 nmol/mL) of Itacitinib (E) or Deucravacitinib (F) followed by stimulation with Art- (50 ng/mL), Phy- (10 ng/mL), or Chi IFN-β (500 ng/mL) for 30 min. Western blots are representative of three independent experiments. (G-H) Flow cytometry ran on AjKi2 cells pretreated with Art IFN-β, Phy IFN-β, or Chi IFN-β or U-IFN then challenged with VSV-GFP (G) or EBOV-GFP (H) after 20 hours. Inhibitory concentration 50 (IC50) values were calculated for each IFN-I. Presented data are representative of three independent experiments conducted in triplicate.

### Art-, Phy-, and Chi IFN-β induce type-I IFN signaling in Jamaican fruit bat cells

To evaluate whether the family and order-level consensus IFN-βs induce IFN-I signaling comparable to conspecific IFN-β in JFB cells, equivalent doses of Art, Phy, or Chi IFN-β were added to primary kidney cells (AjKi2). The IFN-I signaling in response to U-IFN was also assessed because it is commercially available and commonly applied to experiments involving diverse host species. Cell lysates collected at 30 minutes (min) and 24 hours (h) were analyzed by Western blot to assess activation of the IFN-I signaling pathway as measured by the phosphorylation (p) of STAT1 and STAT2 and the increased expression of ISGs, total STAT1, total STAT2, ISG15, and IFIT3 (Fig. 1B). Following treatment with any of the IFN-Is, the level of pSTAT1 and pSTAT2 increased at 30 min, and the expression of ISGs, STAT2 and ISG15, increased at 24 h (Fig.1C). Increases in total STAT1 and IFIT3 were observed following stimulation with Art IFN-β and Phy IFN-βs, but limited or absent following stimulation with Chi IFN-β or U-IFN (Fig. 1C).

To quantify the results for pSTAT1 and pSTAT2 densitometry analysis was performed. The pixel densities of pSTAT1 and pSTAT2 were normalized to the actin band to account for differences in the total amount of protein across samples. To enable a visual comparison across treatment groups, the actin-normalized pSTAT1 and pSTAT2 values were normalized to Chi IFN-β at 30 min. All four IFN-Is induced the type-I signaling pathway, however, the duration of pSTAT1 and -2 was longer following treatment with Art- and Phy IFN-βs than Chi IFN-β and U-IFN (Fig. 1D).

To validate that all IFN-Is activated the IFN-I signaling pathway as expected, JFB cells were treated with two IFN-I signaling inhibitors. Deucravacitinib and itacitinib inhibit tyrosine kinase 2 (TYK2) and Janus kinase 1 (JAK1), respectively, to prevent STAT1 and STAT2 phosphorylation (Fig. 1B). JFB cells were treated with a dilution series of either inhibitor for 24 h then treated with Art-, Phy-, or Chi IFN-β for 30 min. As expected, pSTAT1 and pSTAT2 signals decreased with increasing itacitinib (Fig. 1E) or deucravacitinib (Fig. 1F) concentration.

After confirming that IFN-I signaling was sensitive to inhibitors, the antiviral activities of the four IFN-Is were compared. JFB cells were treated with Art-IFN-β, Phy-IFN-β, or Chi IFN-β or U-IFN for 24 h, then infected with either recombinant vesicular stomatitis virus (VSV) or Ebola virus (EBOV) expressing green fluorescent protein (GFP) reporter gene for 20 h. The concentration required to decrease the number of GFP-positive cells by 50% (IC50) was determined for each of the IFN-Is using flow cytometry. Phy IFN-β (0.74 ng/mL) inhibited VSV-GFP most effectively, followed by Art IFN-β (7.45 ng/mL), while Chi IFN-β (54.71 ng/mL) and U-IFN (62.09 ng/mL) had similar efficacy (Fig. 1G). When tested for antiviral activity against EBOV-GFP, similar results were observed, with Chi IFN-β (336.6 ng/mL) and U-IFN (324.9 ng/mL) requiring higher inhibitory concentrations compared to Art IFN-β (7.09 ng/mL) and Phy IFN-β (0.47 ng/mL) (Fig. 1H).

### Phy- and Chi IFN-β and U-IFN induce ISG repertoires comparable to conspecific IFN-β

Next-generation RNA-sequencing (RNA-seq) was performed in JFB cells to compare the ISG repertoires induced following treatment with each of the IFN-Is. Prior to submitting samples for RNA-seq, the doses of each IFN-I inducing 80% of the maximum ISG transcription were determined. Consistent with the antiviral assays, lower concentrations of Phy IFN-β (0.27 ng/mL) and Art IFN-β (2.4 ng/mL) were required to induce equivalent transcription of the ISGs *Isg15*, *Mx1*, and *Oas1* compared to Chi IFN-β (973 ng/mL) and U-IFN (762 ng/mL) (Supplementary Fig. 2A-C).

JFB cells were stimulated with functionally equivalent doses of Art IFN-β (n=3), Phy IFN-β (n=3), or Chi IFN-β (n=3) or U-IFN (n=3) or mock (n=3) treated for 20 h prior to RNA collection. Prior to submitting the samples for RNA-seq, host gene RT-qPCR was performed to ensure similar induction of ISGs (Supplementary Fig. 2B). All four IFN-Is induced ISGs within five-fold, supporting that the selected concentrations were appropriate for comparison.

All four IFN-Is induced (log_2_ fold-change ≥ 2, adjusted *p*-value ≤0.05) expression of 147 shared genes (Fig. 2A). Many of these 147 shared genes are described ISGs in multiple host species [28, 29] and have comparable fold induction across the IFN-I treatment groups (Fig. 2D). Most of the genes that were induced significantly in response to treatment with one or more IFN-I, but not all four, trended for induction in all groups and likely were excluded due to the stringent cut-off values or intersample variation (Sup. Fig. 3, Sup. Table 2). In support of the similar ISG repertoires across the four IFN-I treatment groups, Gene Ontology (GO) analysis identified the same groups of biological functions significantly enriched including: type I interferon response, type II interferon response, response to/regulation of virus infection, antigen processing and presentation, regulation of adaptive immune responses, regulation of cytokine production and signaling, negative regulation of immune response, regulation of dendritic cells, and natural killer cell regulation (Fig.2C; Sup. Tables 3-6). The only unique biological function within the top 100 GO IDs for each of the IFN-I treatment groups was inflammasome regulation in Phy IFN-β treated cells (Fig. 2C). The inflammasome GO terms were significantly enriched in the other groups but were not in the top 100 GO IDs (Sup. Tables 3-6). Overall, each of the four IFN-Is induced comparable ISG repertoires.

**Fig 2.**
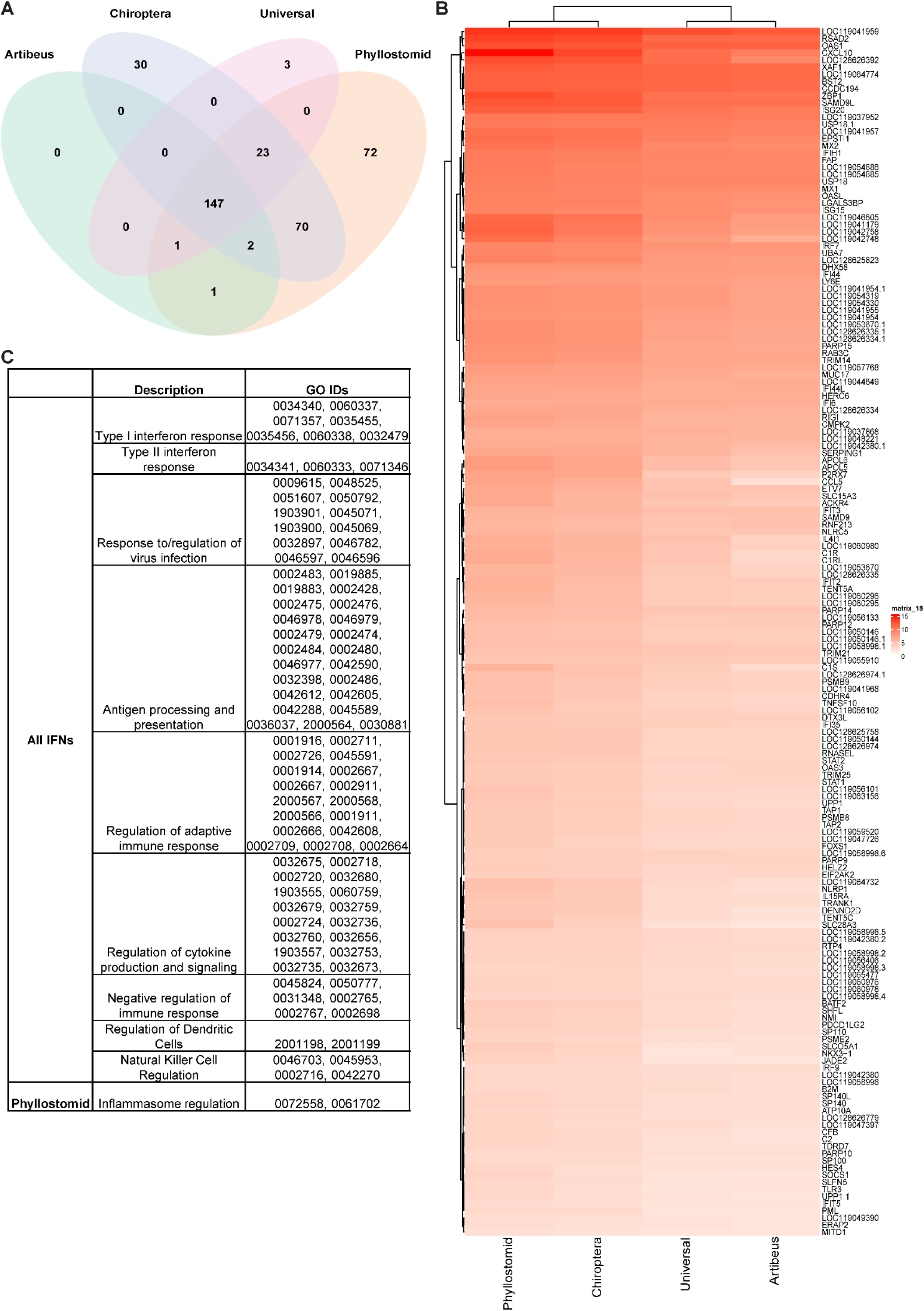
Phyllostomid and Chiroptera consensus IFN-β and U-IFN induce ISG repertoires comparable to conspecificIFN-β. Primary AjKi2 cells were stimulated with Artibeus- (n=3), Phyllostomid- (n=3), or Chiroptera- (n=3), IFN-β or universal IFN (n-3) for 16 hours then their transcriptomes were assessed and compared to mock stimulated cells (n-3). (A) The number of upregulated genes by each of the IFN-I types are presented as a Venn diagram. (B) The shared (n=147) genes upregulated by each of the IFN-I types are presented as a heat map. (C) The top 100 gene ontology (GO) enrichment identification (ID) codes and description of upregulated biological processes for each of the IFN-I types.

### Chi IFN-β induces type-I signaling across diverse bat cell lines

To determine whether Chi IFN-β is effective in other bat cell lines, IFN-I signaling, inhibition, and antiviral assays were performed in bat cell lines from five distinct bat families (Table 1). Each bat cell line was treated with 500 ng/mL Art IFN-β or Chi IFN-β, or U-IFN for 30 min or 24 h to evaluate IFN-I pathway activation (Sup. Fig. 4A-I). Among the Yangochiropteran cell lines, Chi IFN-β induced stronger IFN-I signaling than U-IFN at 30 min in PipNi (Fig. 3A, Sup Fig. 4A), SubK12-08 (Fig. 3B, Sup Fig. 4B), and TbLu1 (Fig. 3C, Sup Fig. 4C). Notably, no signal for pSTAT2 or total STAT2 was detected in the PipNi cell lines, likely due to poor epitope cross-reactivity (Sup. Fig. 4A). The IFN-I signaling pathway was activated in response to both Chi IFN-β and U-IFN in each of the Yinpterochiropteran cell lines (Fig. 3D-I; Sup. Fig. 4D-I). U-IFN induced stronger (30 min) and/or more sustained (24 h) pSTAT1 and pSTAT2 in Pteropodidae cell lines (PaKiT (Fig. 3D, Sup Fig. 4D), HypNi (Fig. 3E, Sup Fig. 4E), RoNi (Fig. 3F, Sup Fig. 4F), EpoNi (Fig. 3G, Sup Fig. 4G), and EidNi (Fig. 3H, Sup Fig. 4H)). In RhiLu cell lines, pSTAT1 and pSTAT2 were similar between Chi IFN-β and U-IFN at both time points (Fig. 3I, Sup Fig 4I). As expected, none of the bat cell lines were responsive to Art IFN-β (Fig. 3A-I; Sup. Fig. 4A-I).

**Table 1.** Description of bat cell lines used to assess Chiroptera IFN-β.

| Bat suborder | Bat family | Species | Cell line | Tissue | Reference |
| --- | --- | --- | --- | --- | --- |
| Yinpterochiroptera | Pteropodidae | *Pteropus alecto* | PaKiT | Kidney | [86] |
|  |  | *Rousettus aegyptiacus* | RoNi | Kidney | [87] |
|  |  | *Eidolon helvum* | EidNi/41.3 | Kidney | [88] |
|  |  | *Epomops buettikoferi* | EpoNi/22.1 | Kidney | [87] |
|  |  | *Hypsignathus monstrosus* | HypNi | Kidney | [87] |
|  | Rhinolophidae | *Rhinolophus alcyone* | RhiLu/1 | Lung | [89] |
| Yangochiroptera | Phyllostomidae | *Artibeus jamaicensis* | AjKi RML2 | Kidney | [19] |
|  | Miniopteridae | *Miniopterus schreibersii* | SubK12_08 | Kidney | [90] |
|  | Molossidae | *Tadarida brasiliensis* | Tb 1 Lu | Lung | ATCC, CCL-88 |
|  | Vespertilionidae | *Pipistrellus pipistrellus* | PipNi | Kidney | [88] |

**Fig 3.**
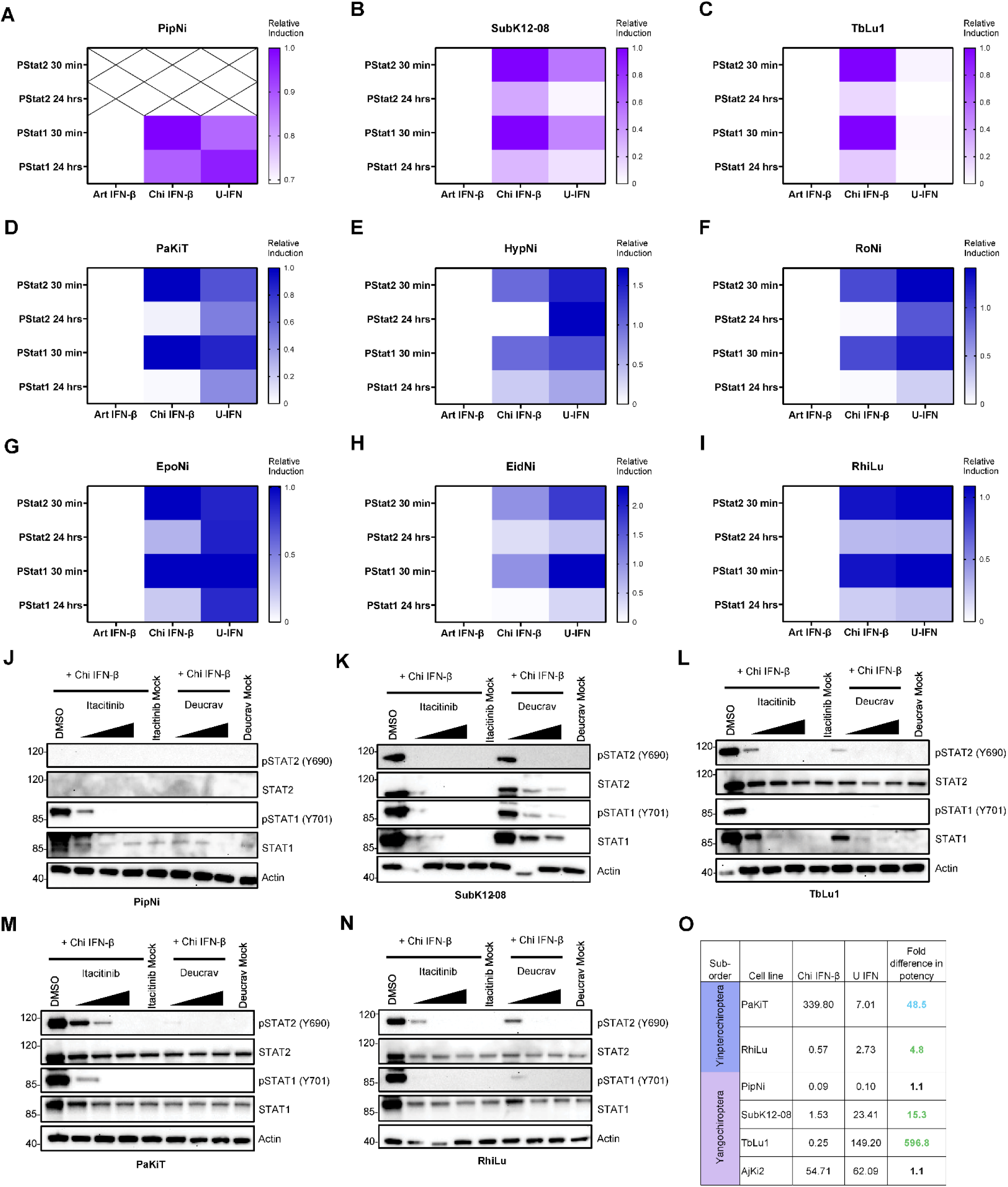
Chiroptera IFN-β induces type-I signaling across diverse bat cell lines. (A-I) Densitometric quantification of immunoblots from bat cell lines belonging to the suborder Yangochiroptera, including PipNi (A) SubK12-08 (B), and TbLu-1 (C), and the suborder Yinpterochiroptera, including PaKiT (D), HypNi (E), RoNi (F), EpoNi(G), EidNi (H), and RhiLu (I). Cells were treated with *Artibeus jamaicensis* (Art), Phyllostomid consensus (Phy), and Chiroptera consensus (Chi) interferon-beta (IFN-β) (500 ng/mL) and lysed at 30 minutes (min) and 24 hours (h). Band intensities were normalized to actin, and then to the Chi IFN-β at 30 min. Representative western blots used for quantification are in Supp Fig 4. (J-N) Immunoblots for PipNi (J), SubK12-08 (K), TbLu-1 (L), PaKiT (M), RhiLu (N) cells treated with three concentrations (1, 5, 25 nmol/mL) of Itacitinib or Deucravacitinib for 24 hours, followed by Chi IFN-β (500 ng/mL) for 30 min. (O) Cells were treated with serial dilutions of the Chi IFN-β or U-IFN for 24 h prior to infection with VSV-GFP and the percentage of GFP positive cells was determined 20 h post-infection. Fold difference in the inhibitory concentration 50 (IC50) dose between Chi IFN-β and U-IFN. Chi IFN-β equally (black), more (green) or less (blue) efficacious than U-IFN. Data used to determine VSV-GFP dose and determine IC50s for each cell line are presented in Sup. Fig. 5.

After confirming activation of the IFN-I response, sensitivity to IFN-I inhibitors and antiviral activity were evaluated in one cell line from each bat family. Only PaKiT cells were used for the IFN-I inhibition and antiviral assays due to the similar phenotype of higher sensitivity to U-IFN observed for all pteropodid cells. Chi IFN-β-induced signaling was responsive to both IFN-I inhibitors, deucravacitinib and itacitinib, in a dose-dependent manner in each of the five bat family representative cell lines: Vespertilionidae (Fig. 3J), Miniopteridae (Fig. 3K), Molossidae (Fig. 3L), Pteropodidae (Fig. 3M), and Rhinolophidae (Fig. 3N). The antiviral activities of Chi IFN-β and U-IFN were compared in cells infected with VSV-GFP (dose determined for each cell line: Sup. Fig. 5A-E). Chi IFN-β was as potent as (PipNi, AjKi2) or more potent (SubK12-08 and TbLu1) than U-IFN in the Yangochiroptera cell lines (Fig. 1G, Fig. 3O, Sup. Fig. 5F-H). RhiLu cells were marginally more sensitive to Chi IFN-β than U-IFN (Fig. 3O, Sup. Fig. 5I), but the PaKiT cells were more sensitive to U-IFN than Chi IFN-β (Fig. 3O, Sup. Fig. 5J). Overall, both Chi IFN-β and U-IFN are antiviral in all bat cell lines evaluated, but their effective concentrations vary.

## Discussion

Comparing the within-host dynamics in response to bat-borne viruses between humans and bats is important for understanding determinants of pathogenesis and the potential to cause human disease. Despite recent advances in resources to study bats and their immune systems – including improvements in the number and quality of bat genomes [30, 31] and induced pluripotent stem cell [32] and organoid development [33, 34] – the resources to dissect complex virus-host interactions in bat cells are limited. The IFN-I pathway is critical in regulating the host response to viral infection. Improved availability of recombinant IFN-Is that are functional in bat cells would bolster the capacity to evaluate viral innate immune antagonism and to compare ISG repertoires and functions. Due to the selective pressures on cytokine-receptor binding interfaces, designing conspecific IFN-Is for each of the 1,500 plus recognized bat species is impractical. Here, we designed and validated a bat order-level consensus IFN-β as a proof of concept that consensus cytokines can be engineered to function across diverse taxa.

IFN-I subtypes include alpha (α), β, epsilon (ε), kappa (κ), and omega (ω). IFN-α, IFN-β, and IFN-ω act broadly, and IFN-ε and IFN-κ are more compartmentalized within mucosal sites and the skin, respectively [35, 36]. For most species, a single *Ifnb1* gene is present within the IFN-I locus, while genes encoding IFN-α and IFN-ω often exist as multiple copies [23]. Evolutionarily, the genes encoding IFN-α and IFN-ω have expanded and contracted in response to different selective pressures which resulted in variation in the sequence and number of genes across mammalian taxa [23, 37]. For example, relative to humans, multiple bat species have expansion of IFN-ω and contraction of IFN-α encoding genes [38]. The diversity in the sequence and number genes precluded the straightforward alignment of IFN-α or IFN-ω encoding genes to generate a consensus. Future studies are needed to evaluate the importance of the different IFN-I subtypes in fine-tuning IFN-I signaling and antiviral responses given the observation that different IFN-I subtypes induce distinct antiviral states and/or ISG repertoires [39–41].

A crucial consideration when developing Chi IFN-β was whether the cellular response would be comparable to conspecific IFN-β. Due to the concern that an order-wide consensus may not be functional, we also designed a family-level consensus as an intermediate between conspecific and Chi IFN-β. JFB cells were treated with Art IFN-β, Phy IFN-β, and Chi IFN-β to evaluate their functional equivalence. All three IFN-βs induced pSTAT1 and pSTAT2 and expression of ISGs, responded to treatment with JAK1 and TYK2 inhibitors, and restricted VSV and EBOV replication. JFB cells were less sensitive to Chi IFN-β than either Art IFN-β or Phy IFN-β, but Chi IFN-β was as effective as the commercially available U-IFN. Unexpectedly, JFB cells appeared more sensitive to Phy IFN-β than Art IFN-β. This phenomenon could be due to Phy IFN-β binding to the IFNAR with a different affinity and/or cause different orientations of the receptor intracellular domains that could alter the kinetics of signal transduction and/or receptor endocytosis [42–46]. Additionally, substantial variation in IFN-β amino acid sequence has been observed across the different published and newly sequences *A. jamaicensis* genomes, and the sequence used for the recombinant IFN-β used in this project was based on the Cold Spring Harbor genome. Thus, within-species variance in IFN-I and IFNAR sequences could influence signaling efficiency.

Cytokines and their receptors co-evolve to optimize binding kinetics, and perturbations to the binding interface could alter the transcriptomic response. The majority (n = 147) of ISGs were induced by all four IFN-Is. The shared upregulated genes include prototypical ISGs that are observed across mammals with characterized functions including: negative feedback of IFN-I signaling (Socs1, Usp18), antigen processing and presentation (B2m, Tap1), antiviral effectors (Bst2, Rnasel), complement (C1r, C1s), chemokines (Ccl5, Cxcl10), pathogen- and damage-recognition receptors (Ifih1, Zbp1), transcription factors (Irf7, Stat2), and regulation of host gene transcription and translation (Pkr, Parp9) [28, 47]. Although the ISG repertoires were similar among the three bat IFN-βs and the commercially available U-IFN, more genes were upregulated in response to Phy IFN-β and Chi IFN-β. The differences in the number of ISGs induced may be explained partially by minor experimental variance in treating the cells with perfectly equivalent doses of the different IFN-Is. In support of this, most of the ISGs that are specific to one or two of the bat IFN-βs are trending toward upregulation but are below the stringent statistical cutoff (Sup. Fig. 3, Sup. Table 2).

Importantly, Chi IFN-β induced IFN-I signaling in bat cells from five additional families. In cell lines from the suborder Yangochiroptera, the Chi IFN-β dose required to inhibit VSV-GFP was either equivalent to (Vespertilionidae, PipNi; Phyllostomidae, AjKi2) or lower (Miniopteridae, SubK12-08; Molossidae, TbLu1) than U-IFN. Within the Yinpterochiroptera cells, Rhinolophid cells (RhiLu) were equally sensitive to Chi IFN-β and U-IFN whereas Pteropodid cells (PaKiT) were nearly 50-fold more sensitive to U-IFN than to Chi IFN-β. The broad range in Chi IFN-β’s antiviral efficacy across the six bat families tested suggests that the consensus sequence may be biased toward Yangochiroptera or away from Pteropodidae or that the *Pteropus alecto* IFN-β/IFNAR interface is more sensitive to the amino acid changes between conspecific- and Chi IFN-β. As expected, only the JFB cells responded to Art IFN-β supporting that species-targeted recombinant IFN-βs are not effective for studying the IFN-I pathway in diverse bat species.

Future studies are necessary to determine the full breadth of Chi IFN-β’s utility. Due to limitations in the diversity of bat cell lines available, we were unable to evaluate Chi IFN-β in the remaining 15 bat families. Most of the bat cell lines that have been generated are from the six families studied here due to their prominence in virus spillover [4]. Further, these six bat families include representatives from both bat suborders supporting the likelihood that Chi IFN-β could be effective for any bat species of interest. Once the Bat1K genome project publishes genomes from each of the 21 bat families, a next-generation consensus Chi IFN-β can be designed that might have balanced efficacy across taxa. An alternative approach to a single, order-wide IFN-β is to manufacture 21 family-level consensus IFN-βs.

Chi IFN-β represents the first engineered order-level consensus IFN-I. The similarity in IFN-I signaling, transcriptomic responses, and antiviral activities between Art- and Chi IFNβ encourages the reliability of Chi IFN-β as a tool to rapidly evaluate innate antiviral immune responses in diverse bat species. This enables the comparison of bat-borne viruses’ antagonism of bat and human immune systems without relying on the over-expression of bat host factors in human cells. This concept can be translated to design other consensus cytokines to accelerate the field’s capacity to study bat immune systems. For example, FLT3L could be generated to differentiate dendritic cells [48] or IL-2 to promote T-cell maturation [49]. Our approach suggests that beyond Chiroptera, cytokines could be designed for additional host of interest at the family or order-level. Overall, this study supports the application of Chi IFN-β to study bats’ IFN-I signaling pathway and to evaluate whether differences in IFN-I signaling or effectors may contribute to bats’ resilience against viruses that are virulent in humans.

## Materials and Methods

### Cells and viruses

Bat cells (Table 1) were maintained in DMEM/F-12 supplemented with 1X MEM non-essential amino acids (NEAA) (Gibco), 10% FBS, 50 U/mL penicillin, and 50 μg/mL streptomycin (10DMEM/F-12). All cells were maintained at 37 *°*C, 5% CO_2_ and were *Mycoplasma* negative. The species of each cell-line was verified by Cytochrome B sequencing. Replication-competent recombinant EBOV [50] and VSV [51] encoding a GFP reporter gene were used for *in vitro* studies. All viruses were sequence verified and confirmed *Mycoplasma* negative. Work with infectious EBOV under Biosafety level 4 (BSL4) conditions was approved by the Institutional Biosafety Committee (IBC) and conducted in RML’s BSL4 facility. Virus inactivation of all samples for flow cytometry was performed according to IBC-approved standard operating procedures (SOPs).

### IFN-β expression and purification

The *Artibeus jamaicensis* (XM_037153906.2) *Ifnb1* sequence was used. The Phy family consensus sequence was designed using the *Ifnb1* gene sequence from all Phyllostomid bat genomes available in the NCBI database as of November 2022 (Sup. Table 1). The Chi *Ifnb1* consensus sequence was design using the *Ifnb1* gene sequences from all bat genomes available in the NCBI database as of November 2022 (Sup. Table 1). To prevent bias from the number of species represented in the bat families, we first generated a consensus of each of the ten bat families used prior to the generation of the Chiroptera consensus. An IL-2 secretion signal, His-tag, and TEV protease signal were added upstream of each *Ifnb1* sequence open reading frame, and the synthesized sequence was inserted into the pcDNA3.4 vector (GenScript). The secretion signal of *Ifnb1* was not used for any of the *Ifnb1* sequences due to substantial variation in the signal sequence which could have impacted expression or secretion efficiency.

Art- and Phy IFN-β were produced and purified as described previously for Chi IFN-β [52]. tsA201 cells were grown in suspension with Gibco® FreeStyle™ 293 Expression Medium plus 2 mM glutamine (FS293EMG) in a shaker flask. For each expression, one frozen vial with 1 × 10^7^ tsA201 cells was inoculated with 40 mL of FS293EMG in a 250 mL flat-bottom cell culture shaker flask. Transfection was started when, cells reached a density of 2 × 10^6^/mL, typically after 3 days. For one liter of culture, 1 mg of pcDNA3.4-*Ifnb1* plasmid (filtered with a 0.22 µm filter) was added to 50 mL of prewarmed Hybridoma-SFM (ThemoFisher, 12045076). Four milliliters of PEI MAX® (Transfection Grade Linear Polyethylenimine Hydrochloride, MW 40,000, Polysciences, 24765-1) at 1 mg/mL was added, then gently mixed to homogenize. After incubation at room temperature for 12–15 min, the mixture was added to the culture, and then the flask was returned to the incubator. After 5 days, the culture was centrifuged at 1000 × g for 15 min using a Sorvall RC3B plus centrifuge. The supernatant was stored at −80 °C for future purification.

The supernatant was filtered through a 0.22 µm filter, then mixed with 5 mL of prewashed HisPur™ Ni-NTA Resin (ThermoFisher Scientific) and rocked overnight at 4 °C. The following day, the supernatant was drained, and the resin was packed into a 5 mL column. AKTA FPLC with Unicorn software (GE Healthcare) was used to wash the column with sodium phosphate buffer, pH 7.0, and 200 mM NaCl at 2 mL/min. The protein was eluted with a linear gradient up to 400 mM imidazole over 30 CV. The IFN-β-containing fractions were pooled (∼28 mL) and injected into a 30 mL Slide-A-Lyze cassette with 10 K MWCO (ThermoFisher Scientific). The cassette was then dialyzed against 1.8 L of 50 mM sodium phosphate buffer, pH 7, three times. After dialysis, the protein concentration was measured using a NanoDrop Microvolume Spectrophotometer (ThermoFisher Scientific). IFN-β was then cleaved by TEV protease to remove the N-terminal His tag. The enzymatic cleavage was completed with 1 mg of TEV per 5 mg of IFN-β and incubated at 4 °C overnight with rocking. The digested mixture was then loaded onto a pre-equilibrated 5 mL Ni-NTA Superflow column (Qiagen) to remove the cleaved His-tag, remaining uncleaved protein, as well as additional impurities, with a flow rate of 1 mL/min. The flowthrough fractions were loaded onto a pre-equilibrated HiTrap Capto Q anion exchange column (Cytiva) to separate IFN-β aggregates from monomer with a flow rate of 2 mL/min. The running buffer was 50 mM sodium phosphate, pH 7, and the gradient was 5 mM/min for NaCl from 0 to 500 mM. IFN-β monomer-containing fractions around 200 mM NaCl were immediately collected and analyzed with SDS-PAGE (Sup. Fig. 1C).

### Genomic DNA extraction

Genomic DNA was extracted from primary AjKi2 cells following the MagAttract HMW DNA kit protocol (Qiagen, 67563). In short, cells were trypsinized, washed in PBS and resuspended in 200 µl PBS before addition of 20 µl Proteinase K to the cells. Subsequently 4 µl RNase A and 150 µl Buffer AL were added before 10-minute incubation at 56°C. Downstream bead purification and wash steps followed the manufactures protocol. Samples were eluted in twice 100 µl dH_2_O. DNA quality was checked for quantity using a Qubit fluorometer (Thermo Fisher) and for purity using a TapeStation (Agilent). One WGS library was sequenced on two PacBio Revio SMRTCells with whole genome libraries using SMRTbell prep kit 3.0. Sample yields 11.37 million ccs read, with mean read length 6.56 kb and mean read quality QV35.1.

### Genome assembly

We used the generated PacBio long reads (SRA: SRR38117686 & SRR38117687) to assemble a set of phased genomes using HifiASM [53], with default parameters including deduplication. Following assembly, we explored if haplotype 1, 2 or the merged genome were the most appropriate to use based on dotplots generated by D-GENIES [54], settling on the combined genome. We also used D-GENIES to align our assembly to the *Artibeus intermedius* (GCA_038363225.1) and *Artibeus lituratus* (GCA_038363175.3) genomes to assign contigs to specific chromosomes. The final assembled genome is available on NCBI (accession: SUB16128800).

We assessed genome assembly and annotation completeness using BUSCO version. 5.0.0 and eukaryote & laurasiatherian OrthoDBs, data version 10 [55]. We assessed the extent of genome contiguity using QUAST [56] compared to the previous *Artibeus jamiciensis* genome assembly.

Our assembled genome is a notable improvement in contiguity on the previous assembly, with several fully assembled chromosomes, and the N50 tripling the value of the previous assembly (Table 2). However, we do find fewer complete BUSCOs, suggesting some data is missing in our assembly.

**Table 2:** Table summarizing the assembly and completeness statistics of the *Artibeus jamaciensis* genome assembled for this study, and the previously assembled genome.

|  | ASSEMBLY 2026 | COLD SPRING HARBOR |
| --- | --- | --- |
| QUAST ANALYSIS |  |  |
| TOTAL LENGTH (MBP) | 2248.18 | 2108.39 |
| LARGEST CONTIG (MBP) | 125.59 | 73.2 |
| # CONTIGS | 268 | 867 |
| # CONTIGS (>= 50KBP) | 148 | 367 |
| N50 (MBP) | 67.87 | 22.09 |
| N90 (MBP) | 18.02 | 3.94 |
| L50 | 12 | 28 |
| L90 | 35 | 112 |
| BUSCO EUKARYOTA |  |  |
| COMPLETE SINGLE-COPY BUSCOS (S) | 249 (97.6%) | 246 (96.4%) |
| COMPLETE DUPLICATED BUSCOS (D) | 4 (1.6%) | 3 (1.2%) |
| FRAGMENTED BUSCOS (F) | 2 (0.8%) | 6 (2.4%) |
| MISSING BUSCOS (M) | 0 (0%) | 0 (0%) |
| BUSCO LAURASIATHERIA |  |  |
| COMPLETE SINGLE-COPY BUSCOS (S) | 11303 (92.4%) | 11977 (97.9%) |
| COMPLETE DUPLICATED BUSCOS (D) | 114 (0.9%) | 73 (0.6%) |
| FRAGMENTED BUSCOS (F) | 128 (1.0%) | 37 (0.3%) |
| MISSING BUSCOS (M) | 689 (5.7%) | 135 (1.1%) |

### Gene and Repeat Annotation

We used RepeatModeler2 [57] to identify repetitive sequences in the *Artibeus jamaicensis* genome assembly. Because several repeats were not classified against the limited internal database of RepeatModeler, we used TransDecoder [58] to generate coding and protein sequences for each repeat sequence. We then used BLASTP (e-value < 0.01) to identify hits in the 2018 release of RepBase [59, 60].

Using this custom library, RepeatMasker [61] was employed to annotate the repetitive portions of the genome (parameters: -u -s -poly -engine rmblast -gff -gccalc), producing a hard-masked version of the genome. Finally, we used the generated GFF and BEDTools [62] to create a soft-masked version of the reference genome.

We find the repeat content of our genome assembly is like those of previous *Artibeus* genome assemblies. Most of the repeat content consists of long interspersed nuclear elements (LINE).

Following assembly, we generated a custom transcriptome using previously generated RNA-sequencing data (SRA: SRR13417529-SRR13417532, SRR2982097-SRR2982108, SRR539297, SRR7910151-SRR7910154) alongside transcriptome data generate for this project (SRA: SRR38117688-SRR38117702). The pipeline first filtered paired-end reads using Cutadapt [63] to remove low-quality sequences and barcodes (parameters: --nextseq-trim=2 --trim-n -n 5 -O 5 -q 10,10 -m 35:35). We then aligned these filtered reads to the assembled genome using Minimap2 [64] in short-read splice-aware mode. Finally, we used Trinity [65] to generate the custom transcriptome using default parameters.

Using the transcriptome generated by Trinity, alongside protein databases from the previously published *Artibeus jamaicensis* genome (GCA_021234435.1), we searched for evidence of genic regions in the soft-masked genome assembly using MAKER2 [66]. We utilized identified repeat regions to ensure repetitive areas were not annotated as genes. After the initial MAKER2 [66] run, we extracted all gene models with a predicted protein length of at least 50 amino acids and an Annotation Edit Distance (AED) score [67] of no more than 0.25 to form a training set for SNAP [68]. The resulting HMM file served as input for the second round of the annotation pipeline, combined with GFF files containing transcript, protein, and repeat evidence from round one. We also concurrently annotated the genome using Helixer [69] via the Galaxy web portal [70]. We merged these two annotations using gffcompare [71] to generate a combined gene annotation. Finally, we functionally annotated the genes using BLAST [72], the UniProt database [73], the previous *Artibeus jamaicensis* annotation and EggNOG-mapper [74].

We find 24,554 genes, with 41,234 transcripts, of which 89% overlaps with the previous genome assembly and 84% are expressed to a reasonable level in at least one RNA-sequencing dataset (TPM > 1) (Table 3).

**Table 3:** Table listing the number of genes annotated in this *Artibeus jamaicensis* genome, if they are highly expressed, functionally annotated and found in the previous assemblies’ annotation.

| Category | Gene | Transcript |
| --- | --- | --- |
| Total | 24554 | 41234 |
| Expressed (> 1 TPM in at least one sample). | 20676 | 35486 |
| Functionally Annotated (eggNOG) | 20580 | 35320 |
| Ortholog in Previous Assembly | 21928 | 38060 |
| Overlap | 18489 | 33814 |

### Gene Orthology Inference

Orthologs were identified using OrthoFinder [75], comparing the proteomes of 26 other bat species outlined in Sup. Table 7. As the *Artibeus intermedius* (GCA_038363225.1) and *Artibeus lituratus* (GCA_038363175.3) genomes available on NCBI lacked any annotation, we crudely annotated the genomes using Helixer [69] on Galaxy [70] (Table 4). We extracted protein sequences from the three *Artibeus* genome annotations using GFFread [71]. Using these protein sets, and those described in Sup. Table 8, we then used OrthoFinder to generate a species tree based on single-copy orthologous genes, which was visualized using APE in R [76]. We find that most of the genes annotated in the *Artibeus* genomes are found across the bat phylogeny, supporting the validity of our genome assembly and annotation (Table 3).

**Table 4:** Genome annotation statistics from the annotations of each *Artibeus* genome annotated in this study.

| Category | *Artibeus jamaicensis* | *Artibeus intermedius* | *Artibeus lituratus* |
| --- | --- | --- | --- |
| Gene | 24554 | 18150 | 18252 |
| Transcript | 41234 | 18150 | 18252 |
| Exon | 412124 | 229071 | 229897 |
| CDS | 388643 | 214206 | 214870 |
| five_prime_UTR | 30522 | 28713 | 29016 |
| three_prime_UTR | 21644 | 20380 | 20411 |

### Western blot analysis

Cell monolayers were lysed in Laemmli buffer (Bio-Rad, 1610737) with 5% beta-mercaptoethanol and boiled for 10 min at 95°C. Lysates were run on 4–12% Bis-Tris NuPAGE gels (Invitrogen) at 150 V for 1 h 15 min then transferred onto methanol-activated PVDF membrane (BioRad) for 1h at 60V. Following a 1 h block in 5% powdered milk, the membranes were washed in buffer (1X TBS with 0.1% tween 20) three times. Membranes were probed with primary antibody overnight rocking at 4 °C. The next day, blots were washed three times, incubated with an HRP-conjugated secondary antibody for 1 h rocking at room temperature, and washed three times before development. Blots were incubated with a 1:1 ratio of peroxidase and enhancer reagents Clarity Western ECL (BioRad) or SuperSignal West Femto (Thermo Scientific) and developed on an iBright imaging system (Thermo Fisher Scientific). Primary antibodies used: 1:1000 IFIT3 (Invitrogen, PA5-22230), 1:1000 ISG15 - 22D2 (Cell Signaling Technology, 2758), 1:1000 pSTAT1 – Y701 (Cell Signaling Technology, 9167S), 1:1000 pSTAT2 – Y690 (Cell Signaling Technology, 88410S), 1:1000 total STAT1 (Cell Signaling Technology, 14994S), 1:1000 total STAT2 (Cell Signaling Technology, 72604S), 1:1000 β-actin (GeneTex, GTX629630). Secondary antibodies used: 1:10,000 Donkey-anti-rabbit (GE Healthcare, NA934) and 1:10,000 Sheep-anti-mouse (GE Healthcare, NA931). Area under the curve (AUC) for western blot bands was measured with ImageJ. Normalized expression of phosphorylated protein was determined with the following formulas: (AUC pSTAT1/AUC β-actin) or (AUC pSTAT2/AUC β-actin). The uncropped western blot images are provided in the source file.

### RNA extractions and RT-qPCR

AjKi_RML2 monolayers were lysed in 600 µL transferred to 70% ethanol prior to extraction of RNA using the Rneasy PLUS extraction kit according to the manufacturer’s instructions (Qiagen). Cellular RNA was used to measure host genes using RT-qPCR (Sup. Table 9) to validate ISG induction prior to sample submission for RNA sequencing. The RNA was diluted 1:20 in molecular grade water (Invitrogen). Fold change was calculated for host genes by dividing the IFN-treated relative gene expression by the average relative gene expression of the mock-treated cells (2−ΔΔCt).

### Transcriptomic gene expression analysis

For expression analysis, we processed Illumina short-read RNA-sequencing data using the RNAseek pipeline (https://github.com/OpenOmics/RNA-seek). The pipeline included quality control via FastQC [77], Preseq, Picard, FastQ Screen [78], Kraken2 [79], QualiMap [80], and RSeQC [81]. The pipeline trims adapters and low-quality reads with Cutadapt [63]. The pipeline then aligns trimmed reads to the hybrid-concatenated genome using the splicing-aware aligner STAR [82] in two-pass mode, mapping against both the genome and the *Artibeus jamaicensis* transcriptome. Finally, it quantifies each genes expression using RSEM [83].

To identify differentially expressed genes between Mock infection treatment samples and other treatments, we used DEseq2 [84]. We normalized data following the recommended pipeline, removing genes with less than 1CPM expression in at least 3 samples, and treated each titer of infection separately. We considered genes with an FDR-adjusted p-value < 0.05 and an absolute log2(Fold-Change) > 2 to be differentially expressed. We used clusterProfiler to identify functional enrichments among differentially expressed genes [85].

### IFN-I signaling

Bat cells were plated at 250,000 cells/mL and incubated at 5% CO_2_, 37°C overnight. Art IFN-β, Phy IFN-β (AjKi2 only), Chi IFN-β, or U-IFN (pbl assay science, 11200-2) were diluted in OptiMEM (Gibco, 31985070) and 100 µL was added to the cells with a final concentration of 500 ng/mL. Cells were incubated with the different IFN-Is for 30 min or 24 h prior to lysis in Laemmli buffer for western blot analysis as described above.

### IFN-I pathway inhibition

Bat cells were treated with 3, 5-fold serial dilutions (25 nmol/mL, 5 nmol/mL, 1 nmol/mL) of IFN-I signaling (Deucravacitinib (Selleckchem, S8879) and Itacitinib (Selleckchem, S7812)) inhibitors or dimethyl sulfoxide (DMSO) vehicle control overnight prior to stimulation with 50 ng/mL Art IFN- β (AjKi2 only), 10 ng/mL Phy IFN-β (AjKi2 only), or 500 ng/mL Ch IFN-β for 30 min. Cell lysates were collected as described above for western blot analysis.

### Anti-viral assay

To assess the anti-viral effect of the different IFN-Is, bat cells were plated overnight at 250,000 cells/mL in 96-well plates and treated with the indicated serial dilutions of IFN-Is (Supplemental Figure X) for at least 20 hours before virus infection. VSV-GFP or EBOV-GFP were diluted in DMEM (Gibco) supplemented with 2% FBS, 1X NEAA and added to cells for 20 hours. After infection, medium was removed and cells were washed once with 1X DPBS prior to the addition of trypsin (2.5% Trypsin, Gibco) for 5-15 min at 37 *°*C, 5% CO_2_. Trypsinized cells were fixed 1:1 in 10% Neutral Formalin Buffer (Cancer Diagnostics Inc) and analyzed by flow cytometry.

### Flow cytometry

Fixed cells were assessed for GFP positivity using a BD FACS Symphony A5 flow cytometer (BD Biosciences). Raw data was exported to FlowJo v10.10 for analysis (BD Biosciences). Debris and dead cells were gated out using a SSC-A and FSC-A gate, followed by doublets gated out with a SSC-A and SSC-H gate. GFP was then gated using an FSC-A and FIT-C gate to characterize the percent of single cells positive for GFP (Sup. Fig. 6)

## Acknowledgements

We thank Marcel Müller for providing the RoNi, EidNi/41.3, EpoNi/22.1, HypNi, RhiLu/1, and PipNi cells, Michelle Baker for providing the PaKiT cells, and Ayato Takada for providing the SubK12_08. We would also like to thank AJ Kniazeff, D Constantine, WA Nelson-Rees, D Schmidt for making the Tb 1 Lu cells available through ATCC.

## Author contributions

Conceptualization: SvT

Investigation: JRB, BB, AW, AM, AC, YH,

Data analysis: TH, LD, SP, PB, CM

Supervision: AW, SvT

Writing—original draft: JRB, BB, SvT

Writing—review & editing: JRB, BB, AW, AM, AC, YH, TH, LD, SP, PB, CM, SvT

## Dava availability

We have summarized the previously published data used in this study in Sup. Table 7. The data generated in this study have been deposited in Figshare at 10.6084/m9.figshare.34613031.

All data generated for this study is available under the NCBI BioProject ID PRJNA1454049. RNA-sequencing are available on the sequence read archive (SRA) under the accessions SRR38117686-SRR38117693, SRR38117696-SRR38117702. PacBio DNA-sequencing are available on the sequence read archive (SRA) under the accessions SRR38117694 & SRR38117695.

## References

1. Simmons, N.B., Cirranello, A.L., Bat Species of the World: A taxonomic and geographic database. 2026.

2. Lei, M. and D. Dong, Phylogenomic analyses of bat subordinal relationships based on transcriptome data. Scientific reports, 2016. 6: p. 27726.

3. Santana, S.E., Bernal-Rivera, A., Chhaya, V., Guerra, E.B., Quinche, L.L., Visconti, F, Law, C.J., An Integrative Perspective on Bat Evolution. Annual Review of Ecology, Evolution, and Systematics, 2025. 56: p. 291–314.

4. Letko, M., et al., Bat-borne virus diversity, spillover and emergence. Nat Rev Microbiol, 2020. 18(1740-1534 (Electronic)).

5. Lytras, S., et al., The animal origin of SARS-CoV-2. Science, 2021. 373(6558): p. 968–970.

6. Meza, D.K., et al., Ecological determinants of rabies virus dynamics in vampire bats and spillover to livestock. Proc Biol Sci, 2022. 289(1471-2954 (Electronic)).

7. Towner, J.S., et al., Marburg virus infection detected in a common African bat. PloS one, 2007. 2(8): p. e764.

8. Towner, J.S., et al., Isolation of genetically diverse Marburg viruses from Egyptian fruit bats. PLoS pathogens, 2009. 5(7): p. e1000536.

9. Chua, K.B., et al., Isolation of Nipah virus from Malaysian Island flying-foxes. Microbes and infection, 2002. 4(2).

10. Anderson, D.E., et al., Isolation and Full-Genome Characterization of Nipah Viruses from Bats, Bangladesh. Emerging infectious diseases, 2019. 25(1).

11. Williamson, M.M., et al., Transmission studies of Hendra virus (equine morbillivirus) in fruit bats, horses and cats. Australian veterinary journal, 1998. 76(12).

12. Williamson, M.M., et al., Experimental hendra virus infection in pregnant guinea-pigs and fruit Bats (Pteropus poliocephalus). J Comp Pathol, 2000. 122(2-3): p. 201–7.

13. Middleton, D.J., et al., Experimental Nipah virus infection in pteropid bats (Pteropus poliocephalus). J Comp Pathol, 2007. 136(4): p. 266–72.

14. Halpin, K., et al., Pteropid bats are confirmed as the reservoir hosts of henipaviruses: a comprehensive experimental study of virus transmission. Am J Trop Med Hyg, 2011. 85(5): p. 946–51.

15. Amman, B.R., et al., Oral shedding of Marburg virus in experimentally infected Egyptian fruit bats (Rousettus aegyptiacus). Journal of wildlife diseases, 2015. 51(1): p. 113–24.

16. Jones, M.E., et al., Experimental Inoculation of Egyptian Rousette Bats (Rousettus aegyptiacus) with Viruses of the Ebolavirus and Marburgvirus Genera. Viruses, 2015. 7(7): p. 3420–42.

17. Ciminski, K., Ran, W., Gorka, M., Lee, J., Malmlov, A., Schinköthe, J., Eckley, M., Murrieta, R.A., Aboellail, T.A., Campbell, C.L., Ebel, G.D., Ma, J., Pohlmann, A., Franzke, K., Ulrich, R., Hoffmann, D., García-Sastre, A., Ma, W., Schountz, T., Beer, M., Schwemmle, M., Bat influenza viruses transmit among bats but are poorly adapted to non-bat species. Nat Microbiol, 2019. 4(12): p. 2298–2309.

18. van Tol, S., Port, J. R., Fischer, R. J., Gallogoly, S., Bushmaker, T., Griffin, A., Schulz, J. E., Carmody, A., Myers, L., Crowley, D. E., Falvo, C. A., Riopelle, J. C., Wickenhagen, A., Clancy, C., Lovaglio, J., Shaia, C., Saturday, G., Prado-Smith, J., He, Y., Lack, J., Martens, C., Anzick, S. L., Kendall, L., Schountz, T., Plowright, R.K., Marzi, A., Munster, V. J., Jamaican fruit bats’ (Artibeus jamaicensis) competence for Ebola virus but not Marburg virus is driven by intrinsic differences in viral entry and IFN-I signaling antagonism. bioRxiv, 2024.

19. Port, J.R., Riopelle, J.C., van Tol, S., Wickenhagen, A., Bohrnsen, E., Sturdevant, D.E., Rosenke, R., Lovaglio, J., Lack, J., Anzick, S.L., Cordova, K., Yinda, K.C., Hanley, P.W., Schountz, T., Kendall, L.V., Shaia, C.I., Saturday, G., Martens, C., Schwarz, B., Munster, V.J., Jamaican fruit bat (Artibeus jamaicensis) insusceptibility to mucosal inoculation with SARS-CoV-2 Delta variant is not caused by receptor compatibility. npj viruses, 2024. 2.

20. Yoneyama, M., et al., The RNA helicase RIG-I has an essential function in double-stranded RNA-induced innate antiviral responses. Nat Immunol, 2004. 5(7): p. 730–7.

21. Kato, H., et al., Differential roles of MDA5 and RIG-I helicases in the recognition of RNA viruses. Nature, 2006. 441(7089): p. 101–5.

22. Boehmer, D. and I. Zanoni, Interferons in health and disease. Cell, 2025. 188(1097-4172 (Electronic)).

23. Xu, L., L. Yang, and W. Liu, Distinct evolution process among type I interferon in mammals. Protein Cell, 2013. 4(1674–8018 (Electronic)).

24. Rehberg, E., et al., Specific molecular activities of recombinant and hybrid leukocyte interferons. J Biol Chem, 1982. 257(0021–9258 (Print)).

25. van Tol, S., Hage, A., Rajsbaum, R., Freiberg, A.N., Pteropus vampyrus TRIM40 Is an Interferon-Stimulated Gene That Antagonizes RIG-I-like Receptors. Viruses, 2023. 15(11).

26. Virtue, E.R., et al., Interferon production and signaling pathways are antagonized during henipavirus infection of fruit bat cell lines. PloS one, 2011. 6(1932–6203 (Electronic)).

27. Friedrichs, V., et al., Selection and stability validation of reference gene candidates for transcriptional analysis in Rousettus aegyptiacus. Sci Rep, 2021. 11(2045–2322 (Electronic)).

28. Shaw, A.E., et al., Fundamental properties of the mammalian innate immune system revealed by multispecies comparison of type I interferon responses. PLoS Biology, 2017. 15(1545-7885 (Electronic)).

29. Gonzalez, V., et al., Bat-specific adaptations in interferon signaling and GBP1 contribute to enhanced antiviral capacity. Nat Commun, 2025. 16(2041-1723 (Electronic)).

30. Jebb, D., et al., Six reference-quality genomes reveal evolution of bat adaptations. Nature, 2020. 583(7817).

31. Teeling, E.C., et al., Bat Biology, Genomes, and the Bat1K Project: To Generate Chromosome-Level Genomes for All Living Bat Species. Annu Rev Anim Biosci, 2018. 6(2165-8110 (Electronic)): p. 23–46.

32. Déjosez, M., et al., Bat pluripotent stem cells reveal unusual entanglement between host and viruses. Cell, 2023. 186(1097-4172 (Electronic)).

33. Kim, H., et al., Diverse bat organoids provide pathophysiological models for zoonotic viruses. Science, 2025. 388(1095-9203 (Electronic)).

34. Kellner, M.J., et al., Bat organoids reveal antiviral responses at epithelial surfaces. Nat Immunol, 2025. 26(1529-2916 (Electronic)).

35. Gibbert, K., et al., IFN-α subtypes: distinct biological activities in anti-viral therapy. Br J Pharmacol, 2013. 168(1476-5381 (Electronic)).

36. Li, S.F., et al., Type I Interferons: Distinct Biological Activities and Current Applications for Viral Infection. Cell Physiol Biochem, 2018. 51(1421-9778 (Electronic)).

37. Shields, L.E., et al., Cross-Species Genome-Wide Analysis Reveals Molecular and Functional Diversity of the Unconventional Interferon-ω Subtype. Front Immunol, 2019. 10(1664-3224 (Electronic)).

38. Zhou, P., et al., Contraction of the type I IFN locus and unusual constitutive expression of IFN-α in bats. Proc Natl Acad Sci U S A, 2016. 113(1091-6490 (Electronic)).

39. Schuhenn, J., et al., Differential interferon-α subtype induced immune signatures are associated with suppression of SARS-CoV-2 infection. Proceedings of the National Academy of Sciences of the United States of America, 2022. 119(8).

40. Moll, H.P., et al., The differential activity of interferon-α subtypes is consistent among distinct target genes and cell types. Cytokine, 2011. 53(1096-0023 (Electronic)).

41. Dickow, J., et al., Diverse Immunomodulatory Effects of Individual IFNα Subtypes on Virus-Specific CD8(+) T Cell Responses. Front Immunol, 2019. 10(1664-3224 (Electronic)).

42. Ng, C.T., et al., Alpha and Beta Type 1 Interferon Signaling: Passage for Diverse Biologic Outcomes. Cell, 2016. 164(1097-4172 (Electronic)).

43. Piehler, J., et al., Structural and dynamic determinants of type I interferon receptor assembly and their functional interpretation. Immunol Rev, 2012. 250(1600-065X (Electronic)).

44. Thomas, C., et al., Structural linkage between ligand discrimination and receptor activation by type I interferons. Cell, 2011. 146(1097-4172 (Electronic)).

45. Sharma, N., G. Longjam, and G. Schreiber, Type I Interferon Signaling Is Decoupled from Specific Receptor Orientation through Lenient Requirements of the Transmembrane Domain. J Biol Chem, 2016. 291(1083-351X (Electronic)).

46. Chmiest, D., et al., Spatiotemporal control of interferon-induced JAK/STAT signalling and gene transcription by the retromer complex. Nat Commun, 2016. 7(2041-1723 (Electronic)).

47. Schneider, W.M., M.D. Chevillotte, and C.M. Rice, Interferon-stimulated genes: a complex web of host defenses. Annu Rev Immunol, 2014. 32(1545-3278 (Electronic)).

48. Laouar, Y., et al., STAT3 is required for Flt3L-dependent dendritic cell differentiation. Immunity, 2003. 19(1074-7613 (Print)).

49. Shouse, A.N., K.M. LaPorte, and T.R. Malek, Interleukin-2 signaling in the regulation of T cell biology in autoimmunity and cancer. Immunity, 2024. 57(1097-4180 (Electronic)).

50. Ebihara, H., et al., In vitro and in vivo characterization of recombinant Ebola viruses expressing enhanced green fluorescent protein. J Infect Dis, 2007. 196: p. S313–22.

51. Marzi, A., et al., Vesicular stomatitis virus-based Ebola vaccines with improved cross-protective efficacy. The Journal of infectious diseases, 2011. 204 **Suppl 3**(Suppl 3): p. S1066–74.

52. van Tol, S., Port, J.R., Fischer, R.J., Gallogly, S., Bushmaker, T., Griffin, A., Schulz, J.E., Carmody, A., Myers, L., Crowley, D.E., Falvo, C.A., Riopelle, J.C., Wickenhagen, A., Clancy, C., Lovaglio, J., Shaia, C., Saturday, G., Prado-Smith, J., He, Y., Lack, J., Martens, C., Anzick, S.L., Kendall, L.V., Schountz, T., Plowright, R.K., Marzi, A., Munster, V.J., Jamaican fruit bats’ competence for Ebola but not Marburg virus is driven by intrinsic differences. Nat Commun, 2025. 16(1).

53. Cheng, H., et al., Haplotype-resolved de novo assembly using phased assembly graphs with hifiasm. Nature methods, 2021. 18(2): p. 170–175.

54. Cabanettes, F. and C. Klopp, D-GENIES: dot plot large genomes in an interactive, efficient and simple way. PeerJ, 2018. 6: p. e4958.

55. Simão, F.A., et al., BUSCO: assessing genome assembly and annotation completeness with single-copy orthologs. Bioinformatics, 2015. 31(19): p. 3210–3212.

56. Gurevich, A., et al., QUAST: quality assessment tool for genome assemblies. Bioinformatics, 2013. 29(8): p. 1072–1075.

57. Flynn, J.M., et al., RepeatModeler2 for automated genomic discovery of transposable element families. Proceedings of the National Academy of Sciences, 2020. 117(17): p. 9451–9457.

58. Haas, B.J., et al., De novo transcript sequence reconstruction from RNA-seq using the Trinity platform for reference generation and analysis. Nature protocols, 2013. 8(8): p. 1494–1512.

59. Jurka, J., et al., Repbase Update, a database of eukaryotic repetitive elements. Cytogenetic and genome research, 2005. 110(1-4): p. 462–467.

60. Kapitonov, V.V. and J. Jurka, A universal classification of eukaryotic transposable elements implemented in Repbase. Nature Reviews Genetics, 2008. 9(5): p. 411–412.

61. Nishimura, D., RepeatMasker. Biotech Software & Internet Report, 2000. 1(1-2): p. 36–39.

62. Quinlan, A.R. and I.M. Hall, BEDTools: a flexible suite of utilities for comparing genomic features. Bioinformatics, 2010. 26(6): p. 841–842.

63. Martin, M., Cutadapt removes adapter sequences from high-throughput sequencing reads. EMBnet. journal, 2011. 17(1): p. 10–12.

64. Li, H., Minimap2: pairwise alignment for nucleotide sequences. Bioinformatics, 2018. 34(18): p. 3094–3100.

65. Grabherr, M.G., et al., Trinity: reconstructing a full-length transcriptome without a genome from RNA-Seq data. Nature biotechnology, 2011. 29(7): p. 644.

66. Holt, C. and M. Yandell, MAKER2: an annotation pipeline and genome-database management tool for second-generation genome projects. BMC bioinformatics, 2011. 12(1): p. 491.

67. Eilbeck, K., et al., Quantitative measures for the management and comparison of annotated genomes. BMC Bioinformatics, 2009. 10(1): p. 67.

68. Korf, I., Gene finding in novel genomes. BMC bioinformatics, 2004. 5(1): p. 59.

69. Holst, F., et al., Helixer–de novo prediction of primary eukaryotic gene models combining deep learning and a hidden Markov model. BioRxiv, 2023: p. 2023.02. 06.527280.

70. Afgan, E., et al., The Galaxy platform for accessible, reproducible and collaborative biomedical analyses: 2018 update. Nucleic acids research, 2018. 46(W1): p. W537–W544.

71. Pertea, G. and M. Pertea, GFF utilities: GffRead and GffCompare. F1000Research, 2020. 9: p. ISCB Comm J–304.

72. Madden, T., The BLAST sequence analysis tool. The NCBI handbook, 2013. 2(5): p. 425–436.

73. Consortium, U., UniProt: a hub for protein information. Nucleic acids research, 2015. 43(D1): p. D204–D212.

74. Huerta-Cepas, J., et al., eggNOG 5.0: a hierarchical, functionally and phylogenetically annotated orthology resource based on 5090 organisms and 2502 viruses. Nucleic acids research, 2019. 47(D1): p. D309–D314.

75. Emms, D.M. and S. Kelly, OrthoFinder: phylogenetic orthology inference for comparative genomics. Genome biology, 2019. 20(1): p. 238.

76. Paradis, E., J. Claude, and K. Strimmer, APE: analyses of phylogenetics and evolution in R language. Bioinformatics, 2004. 20(2): p. 289–290.

77. Bioinformatics, B., FastQC. 2023.

78. Wingett, S.W. and S. Andrews, FastQ Screen: A tool for multi-genome mapping and quality control. F1000Res, 2018. 7: p. 1338.

79. Wood, D.E., J. Lu, and B. Langmead, Improved metagenomic analysis with Kraken 2. Genome biology, 2019. 20(1): p. 257.

80. García-Alcalde, F., et al., Qualimap: evaluating next-generation sequencing alignment data. Bioinformatics, 2012. 28(20): p. 2678–2679.

81. Wang, L., S. Wang, and W. Li, RSeQC: quality control of RNA-seq experiments. Bioinformatics, 2012. 28(16): p. 2184–2185.

82. Dobin, A., et al., STAR: ultrafast universal RNA-seq aligner. Bioinformatics, 2013. 29(1): p. 15–21.

83. Li, B. and C.N. Dewey, RSEM: accurate transcript quantification from RNA-Seq data with or without a reference genome. BMC bioinformatics, 2011. 12(1): p. 323.

84. Love, M.I., W. Huber, and S. Anders, Moderated estimation of fold change and dispersion for RNA-seq data with DESeq2. Genome biology, 2014. 15(12): p. 550.

85. Wu, T., et al., clusterProfiler 4.0: A universal enrichment tool for interpreting omics data. The innovation, 2021. 2(3).

86. Crameri, G., et al., Establishment, immortalisation and characterisation of pteropid bat cell lines. PLoS One, 2009. 4(1932-6203 (Electronic)).

87. Kühl, A., et al., Comparative analysis of Ebola virus glycoprotein interactions with human and bat cells. J Infect Dis, 2011. 204(1537-6613 (Electronic)): p. S840–9.

88. Biesold, S.E., et al., Type I interferon reaction to viral infection in interferon-competent, immortalized cell lines from the African fruit bat Eidolon helvum. PloS one, 2011. 6(1932-6203 (Electronic)).

89. Hoffmann, M., et al., Differential sensitivity of bat cells to infection by enveloped RNA viruses: coronaviruses, paramyxoviruses, filoviruses, and influenza viruses. PloS one, 2013. 8(1932-6203 (Electronic)).

90. Maruyama, J., et al., Characterization of the envelope glycoprotein of a novel filovirus, lloviu virus. J Virol, 2014. 88(1098-5514 (Electronic)): p. 99–109.

