## Supplemental Figures for "A Synthetic Consensus Chiroptera IFN-β Induces IFN-I Signaling Across Diverse Bat Taxa"

This file includes:

Supplementary Table 1. Bat species used to design the consensus Chiroptera IFN-β sequence
Supplementary Table 7. List of genomes used in the OrthoFinder analysis
Supplementary Table 9. List of primers and probes for the target genes to evaluate host gene expression changes in *Artibeus jamaicensis* cells
Supplementary Figure 1. Generation of recombinant bat IFN-βs
Supplementary Figure 2. Determination of functionally equivalent doses to compare interferon stimulated gene repertoires
Supplementary Figure 3. Differentially induced genes share similar expression patterns across the IFN-I treatment groups
Supplementary Figure 4. Chiroptera IFN-β induces IFN-I signaling in cell lines from diverse bat taxa
Supplementary Figure 5. Chiroptera IFN-β induces an antiviral state in cell lines from diverse bat taxa
Supplementary Figure 6. Gating strategy to evaluate infection with VSV- or EBOV-GFP

Supplementary Tables Attached as Separate Excel File due to Size:

Supplementary Table 2. Table of differentially expressed genes identified following stimulation with the different IFN-I types
Supplementary Table 3. Gene ontology (GO) terms and descriptions of biological processes differentially regulated following AjKi2 cells stimulated with conspecific IFN-β
Supplementary Table 4. Gene ontology (GO) terms and descriptions of biological processes differentially regulated following AjKi2 cells stimulated with Phyllostomid consensus IFN-β
Supplementary Table 5. Gene ontology (GO) terms and descriptions of biological processes differentially regulated following AjKi2 cells stimulated with consensus Chiroptera IFN-β
Supplementary Table 6. Gene ontology (GO) terms and descriptions of biological processes differentially regulated following AjKi2 cells stimulated with Universal IFN-I
Supplementary Table 8. List of proteins used to generate a species-based tree

**Supplementary Table 1. Bat species used to design the consensus Chiroptera IFN-β sequence**

| Sub-order | Family | Species | Genome Accession |
|---|---|---|---|
| Yangochiroptera | Miniopteridae | Miniopterus natalensis | GCA_001595765.1 |
| | | Miniopterus schreibersii | GCA_004026525.1 |
| | Molossidae | Molossus molossus | GCA_014108415.1 |
| | | Tadarida brasiliensis | GCA_004025005.1 |
| | Mormoopidae | Mormoops blainvillei | GCA_004026545.1 |
| | | Pteronotus mesoamericanus | GCA_021234165.1 |
| | | Pteronotus parnellii | GCA_000465405.1 |
| | Noctilionidae | Noctilio leporinus | GCA_004026585.1 |
| | Phyllostomidae | Anoura caudifer | GCA_004027475.1 |
| | | Artibeus jamaicensis | GCA_021234435.1 |
| | | Carollia perspicillata | GCA_004027735.1 |
| | | Desmodus rotundus | GCA_022682495.2 |
| | | Macrotus californicus | GCA_007922815.1 |
| | | Micronycteris hirsuta | GCA_004026765.1 |
| | | Phyllostomus discolor | GCA_004126475.3 |
| | | Phyllostomus hastatus | GCA_019186645.2 |
| | | Tonatia bakeri | GCA_004024845.1 |
| | Vespertilionidae | Eptesicus fuscus | GCA_000308155.1 |
| | | Ia io | GCA_025583905.1 |
| | | Lasiurus borealis | GCA_004026805.1 |
| | | Lasiurus cinereus | GCA_011751095.1 |
| | | Murina feae | GCA_004026665.1 |
| | | Myotis brandtii | GCA_000412655.1 |
| | | Myotis davidii | GCA_000327345.1 |
| | | Myotis lucifugus | GCA_000147115.1 |
| | | Myotis myotis | GCA_014108235.1 |
| | | Nycticeius humaeralis | GCA_007922795.1 |
| | | Pipistrellus kuhlii | GCA_014108245.1 |
| | | Pipistrellus pipistrellus | GCA_903992545.1 |
| | | Pipistrellus pygmaeus | GCA_004026625.1 |
| Yinpterochiroptera | Pteropodidae | Cynopterus brachyotis | GCA_009793145.1 |
| | | Eidolon helvum | GCA_000465285.1 |
| | | Eonycteris spelaea | GCA_003508835.1 |
| | | Macroglossus sobrinus | GCA_004027375.1 |
| | | Pteropus alecto | GCA_000325575.1 |
| | | Pteropus medius | GCA_902729225.1 |
| | | Pteropus pselaphon | GCA_014363405.1 |
| | | Pteropus vampyrus | GCA_000151845.2 |
| | | Rousettus aegyptiacus | GCA_014176215.2 |
| | | Rousettus leschenaultii | GCA_015472975.1 |
| | Rhinolophidae | Rhinolophus ferrumequinum | GCA_004115265.3 |
| | | Rhinolophus sinicus | GCA_001888835.1 |
| | Hipposideridae | Hipposideros armiger | GCA_001890085.1 |
| | | Hipposideros galeritus | GCA_004027415.1 |
| | | Hipposideros pendleburyi | GCA_021464545.1 |
| | Craseonycteridae | Craseonycteris thonglongyai | GCA_004027555.1 |

**Supplementary Table 7. List of genomes used in the OrthoFinder analysis.** The accession numbers of those genomes, the phylogenetic group these species belong to, and the IDs used in this study.

| Species | Family | Accession | Annotated ID |
|---|---|---|---|
| *Artibeus intermedius* | Phyllostomidae | GCA_038363225.1 | Artibeus_intermedius |
| *Artibeus lituratus* | Phyllostomidae | GCA_038363175.3 | Artibeus_lituratus |
| *Artibeus jamaicensis* | Phyllostomidae | GCA_021234435.1 | Artibeus_jamaicensis_ref |
| *Cnephaeus nilssonii* | Vespertilionidae | GCA_951640355.1 | Cnephaeus_nilssonii |
| *Desmodus rotundus* | Phyllostomidae | GCA_022682495.2 | Desmodus_rotundus |
| *Eptesicus fuscus* | Vespertilionidae | GCA_027574615.1 | Eptesicus_fuscus |
| *Glossophaga mutica* | Phyllostomidae | GCA_039655065.1 | Glossophaga_mutica |
| *Hipposideros armiger* | Hipposideridae | GCA_001890085.1 | Hipposideros_armiger |
| *Hipposideros larvatus* | Hipposideridae | GCA_031876335.1 | Hipposideros_larvatus |
| *Miniopterus natalensis* | Vespertilionidae | GCA_001595765.1 | Miniopterus_natalensis |
| *Miniopterus schreibersii* | Vespertilionidae | GCA_964146895.2 | Miniopterus_schreibersii |
| *Molossus molossus* | Molossidae | GCA_014108415.1 | Molossus_molossus |
| *Molossus nigricans* | Molossidae | GCA_039880945.1 | Molossus_nigricans |
| *Myotis daubentonii* | Vespertilionidae | GCA_963259705.1 | Myotis_daubentonii |
| *Myotis myotis* | Vespertilionidae | GCA_014108235.1 | Myotis_myotis |
| *Phyllostomus discolor* | Phyllostomidae | GCA_004126475.3 | Phyllostomus_discolor |
| *Pipistrellus kuhlii* | Vespertilionidae | GCA_014108245.1 | Pipistrellus_kuhlii |
| *Plecotus auritus* | Plecotus | GCA_963455305.1 | Plecotus_auritus |
| *Pteronotus mesoamericanus* | Mormoopidae | GCA_021234165.1 | Pteronotus_mesoamericanus |
| *Pteropus alecto* | Pteropodidae | GCA_000325575.1 | Pteropus_alecto |
| *Pteropus vampyrus* | Pteropodidae | GCA_000151845.2 | Pteropus_vampyrus |
| *Rhinolophus ferrumequinum* | Rhinolophidae | GCA_004115265.3 | Rhinolophus_ferrumequinum |
| *Rhynchonycteris naso* | Emballonuridae | GCA_037038545.1 | Rhynchonycteris_naso |
| *Rousettus aegyptiacus* | Pteropodidae | GCA_014176215.1 | Rousettus_aegyptiacus |
| *Saccopteryx leptura* | Emballonuridae | GCA_036850995.1 | Saccopteryx_leptura |
| *Sturnira hondurensis* | Phyllostomidae | GCA_014824575.2 | Sturnira_hondurensis |

**Supplementary Table 9. List of primers and probes for the target genes to evaluate host gene expression changes in *Artibeus jamaicensis* cells**

| Primer/probe name | Sequence |
|---|---|
| ISG15 F | CAGAAGGTGGCTGAGCTGAA |
| ISG15 P | FAM-TGGCTGAGTTTCCAGGGGAGGCCC |
| ISG15 R | CTTGTATTCCTTCAGCTGCGC |
| Ifit1 F | AGAGCTTGAAGCAGGCTGAA |
| Ifit1 P | FAM-ACATGCTGGCCAGTCGGAGGTGAG |
| Ifit1 R | CTTCTAGTCTGCCCATGCGG |
| Mx1 F | GTTCTTCATGCTCCGGTCGT |
| Mx1 P | FAM-GCCAGAAGCTGAGCAATGCCATGTTGC |
| Mx1 R | TTCCTCTTGTCGCTGGTGTC |
| OAS1 F | CGTGGTGCAGAGTCACAGAT |
| OAS1 P | FAM-CGCGTGATACGGAAACCTCGCTCGC |
| OAS1 R | GGGCAGGACATCAAACTCCA |
| Ddx58 F | GGAAGAACCCGAATGCCAGA |
| Ddx58 P | FAM-CAGCCCACTGAAGCCCAGAAATTAC |
| Ddx58 R | TTGATAGCAGGAGCAGCGAG |
| HPRT F | AGATGGTGAAGGTCGCAAG |
| HPRT P | FAM-ACTTTGTTGGATTTGAAATTCCAGACAAGTTTG |
| HPRT R | CCTGAAGTATTCATTATAGTCAAGGG |

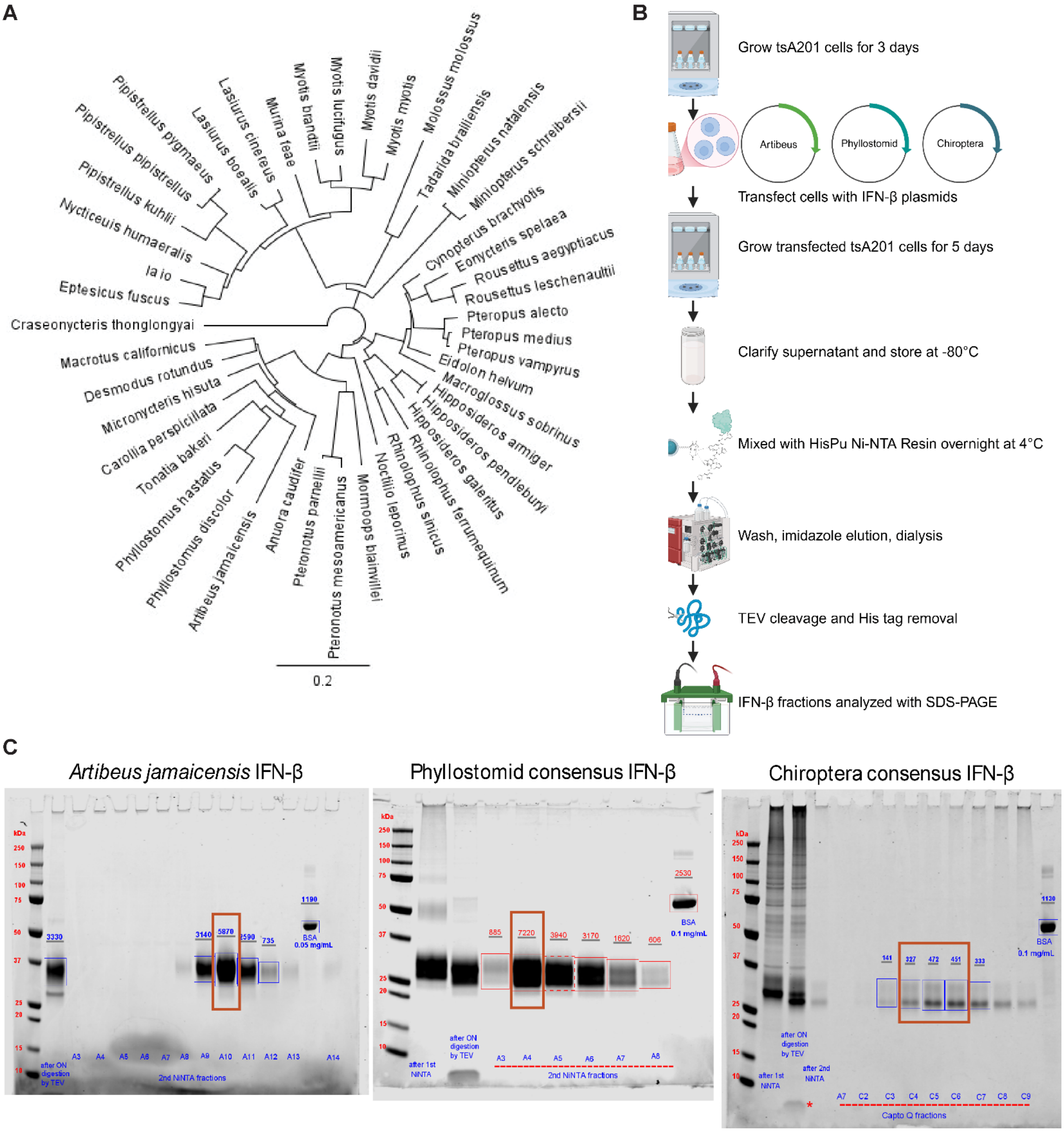

A
Craseonycteris thonglongyai
Pipistrellus pygmaeus
Pipistrellus pipistrellus
Pipistrellus kuhlii
Nycticeius humeralis
Ia io
Eptesicus fuscus
Lasiurus boealis
Lasiurus cinereus
Murina feae
Myotis brandtii
Myotis lucifugus
Myotis davidii
Myotis myotis
Molossus molossus
Tadarida brasiliensis
Miniopterus natalensis
Miniopterus schreibersii
Cynopterus brachyotis
Eonycteris spelaea
Rousettus aegyptiacus
Rousettus leschenaultii
Pteropus alecto
Pteropus medius
Pteropus vampyrus
Eidolon helvum
Macroglossus sobrinus
Hipposideros armiger
Hipposideros pendleburyi
Hipposideros galeritus
Rhinolophus ferrumequinum
Rhinolophus sinicus
Noctilio leporinus
Mormoops blainvillei
Pteronotus mesoamericanus
Pteronotus parnellii
Anuora caudifer
Artibeus jamaicensis
Phyllostomus discolor
Phyllostomus hastatus
Tonatia bakeri
Carollia perspicillata
Micronycteris hisuta
Desmodus rotundus
Macrotus californicus
0.2
B
Grow tsA201 cells for 3 days
Artibeus
Phyllostomid
Chiroptera
Transfect cells with IFN-β plasmids
Grow transfected tsA201 cells for 5 days
Clarify supernatant and store at -80°C
Mixed with HisPu Ni-NTA Resin overnight at 4°C
Wash, imidazole elution, dialysis
TEV cleavage and His tag removal
IFN-β fractions analyzed with SDS-PAGE
C
Artibeus jamaicensis IFN-β
Phyllostomid consensus IFN-β
Chiroptera consensus IFN-β
kDa
250
150
100
75
50
37
25
20
15
10
3330
3140
5870
2590
735
1190
BSA
0.05 mg/mL
after ON digestion by TEV
A3 A4 A5 A6 A7 A8 A9 A10 A11 A12 A13 A14
2nd NiNTA fractions
2530
885
7220
3940
3170
1620
606
BSA
0.1 mg/mL
after 1st NiNTA
after ON digestion by TEV
A3 A4 A5 A6 A7 A8
2nd NiNTA fractions
1130
141
327
472
451
333
BSA
0.1 mg/mL
after ON digestion by TEV
after 1st NiNTA
after 2nd NiNTA
A7 C2 C3 C4 C5 C6 C7 C8 C9
Capto Q fractions

**Supplementary Fig 1. Generation of recombinant bat IFN-βs**

(A) Phylogenetic tree representing the species used to generate the Chiroptera consensus IFN-β, the full list of species and accession numbers for the genomes used is presented in Sup. Table 1. (B) Schematic of the protein synthesis and purification pipeline used to generate the recombinant bat IFN-βs. (C) Protein gel showing the purified IFN-β fractions that were purified. The dark red box indicates the fraction or fractions that were used for each of the recombinant bat IFN-βs.

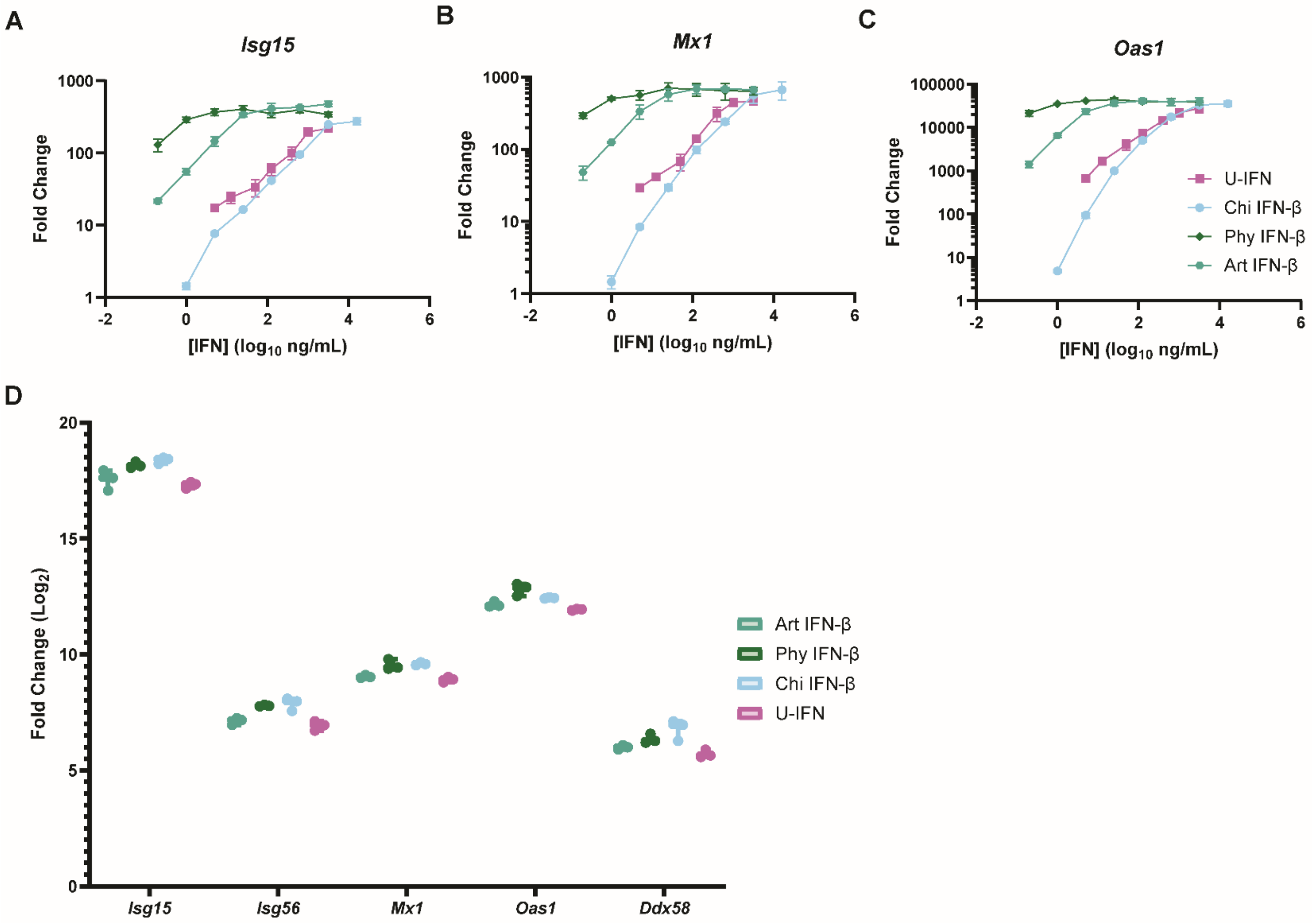


**Supplementary Fig 2. Determination of functionally equivalent doses to compare interferon stimulated gene repertoires**

(A-C) Primary Jamaican fruit bat cells cells were stimulated with serial dilutions of *Artibeus jamaicensis* (Art), Phyllostomid (Phy), or Chiroptera (Chi) interferon-β (IFN- β) or universal type I IFN (U-IFN) for 16 hours then their RNA was collected and evaluated for fold changes of IFN-stimulated genes (ISG), *Isg15* (A), *Mx1* (B), and *Oas1* (C). (D) AjKi2 cells were stimulated with Artibeus- (n=3), Phyllostomid- (n=3), or Chiroptera- (n=3), IFN-β or universal IFN (n-3) for 16 hours then expression of ISGs was assessed and compared to mock stimulated cells (n=3) using RT-qPCR prior to sending the RNA samples for transcriptomics.

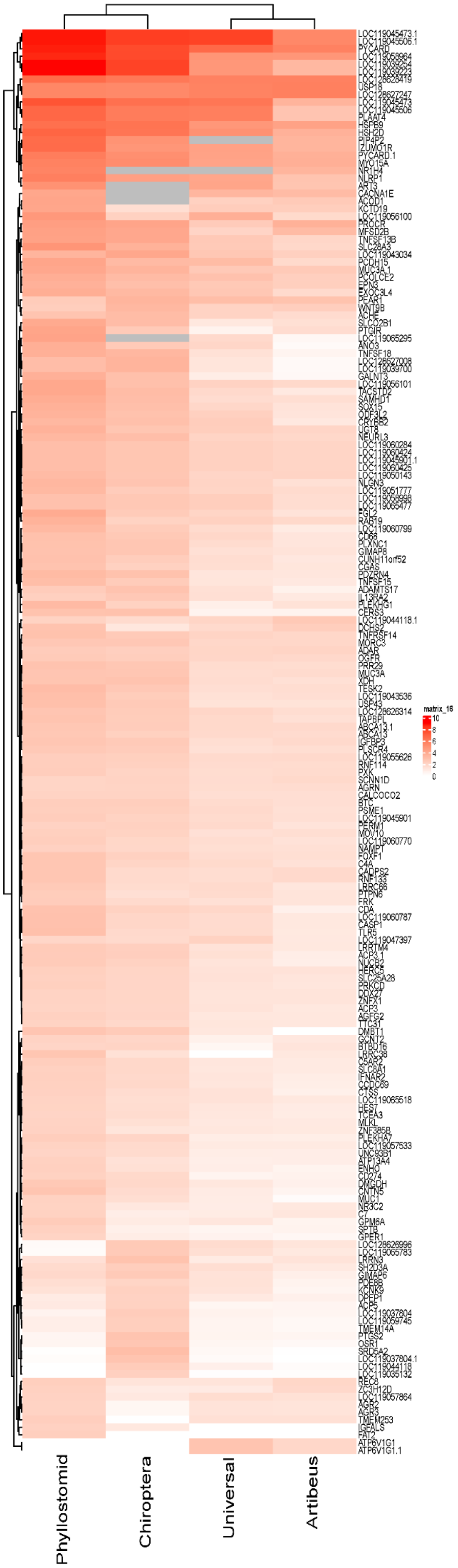
matrix_16
10
8
6
4
2
0
Phyllostomid
Chiroptera
Universal
Artibeus

**Supplementary Fig 3. Differentially induced genes share similar expression patterns across the IFN-I treatment groups**

Primary AjKi2 cells were stimulated with Artibeus- (n=3), Phyllostomid- (n=3), or Chiroptera- (n=3), IFN-β or universal IFN (n-3) for 16 hours then their transcriptomes were assessed and compared to mock stimulated cells (n-3). The genes (n= 202) upregulated by at least one, but not all, IFN-I types are presented as a heat map.

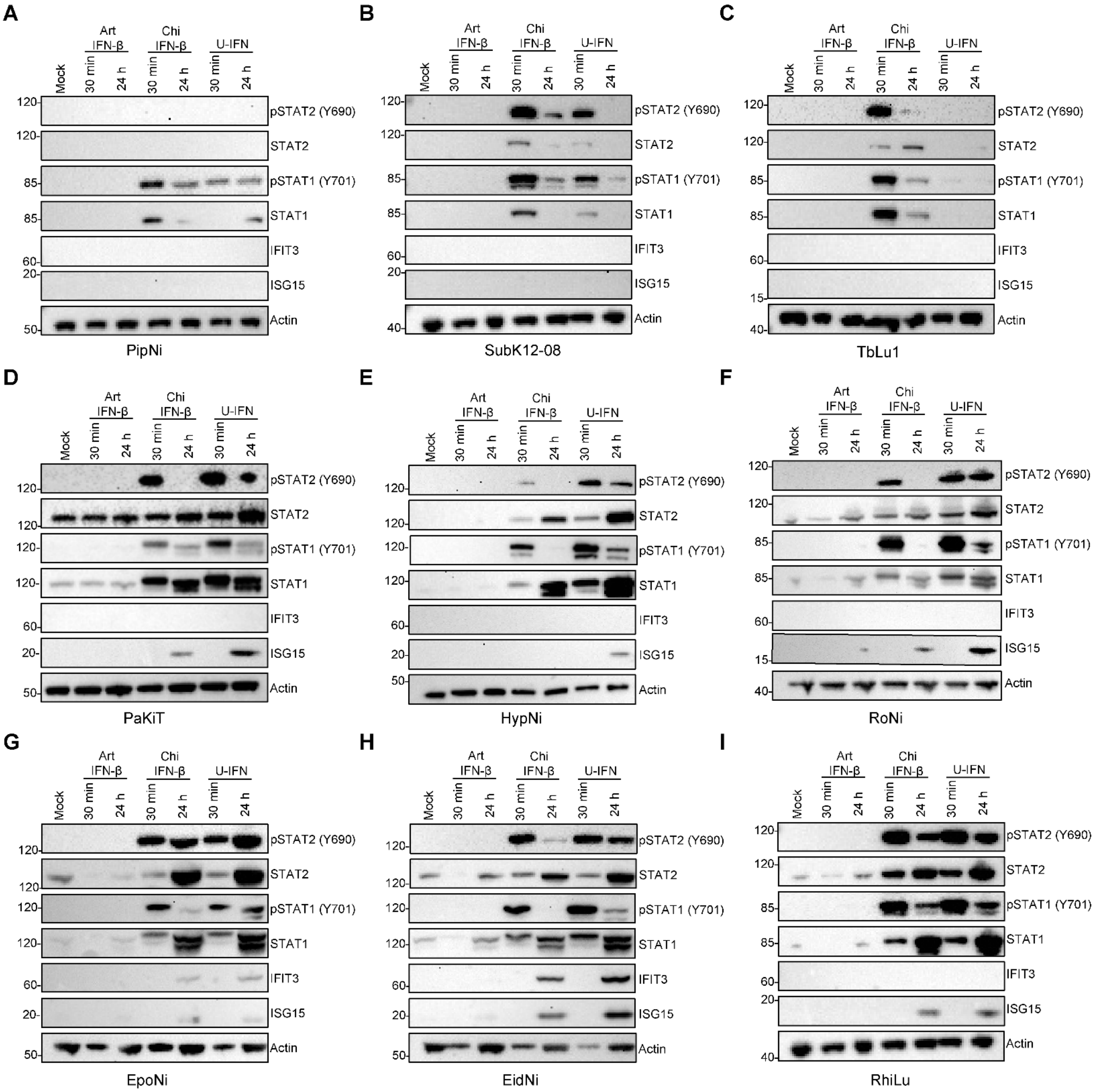
A
Art IFN-β
Chi IFN-β
U-IFN
Mock
30 min
24 h
pSTAT2 (Y690)
STAT2
pSTAT1 (Y701)
STAT1
IFIT3
ISG15
Actin
120
85
60
20
50
PipNi
B
SubK12-08
40
C
TbLu1
15
D
PaKiT
E
HypNi
F
RoNi
G
EpoNi
H
EidNi
I
RhiLu

**Supplementary Fig 4. Chiroptera IFN-β induces IFN-I signaling in cell lines from diverse bat taxa**

Immunoblots of PipNi (A), SubK12-08 (B), TbLu-1 (C), PaKiT (D), HypNi (E), RoNi (F), EpoNi (G), EidNi (H), and RhiLu (I) cells treated with *Artibeus jamaicensis* (Art), Chiroptera (Chi) IFN-β, or Univeral type I IFN (U-IFN) (500 ng/mL) for 30 minutes (min) or 24 hours (h). Blots were probed with phosphorylated (p) STAT1 and STAT2 to evaluate activation of the IFN-I receptor, and STAT1, STAT2, IFIT3, and ISG15 are prototypical IFN-stimulated genes. β-actin was also evaluated as a loading control. Blots shown are representative of at least three independent experiments and were used to calculate densitometry shown in Fig. 3 A-I.

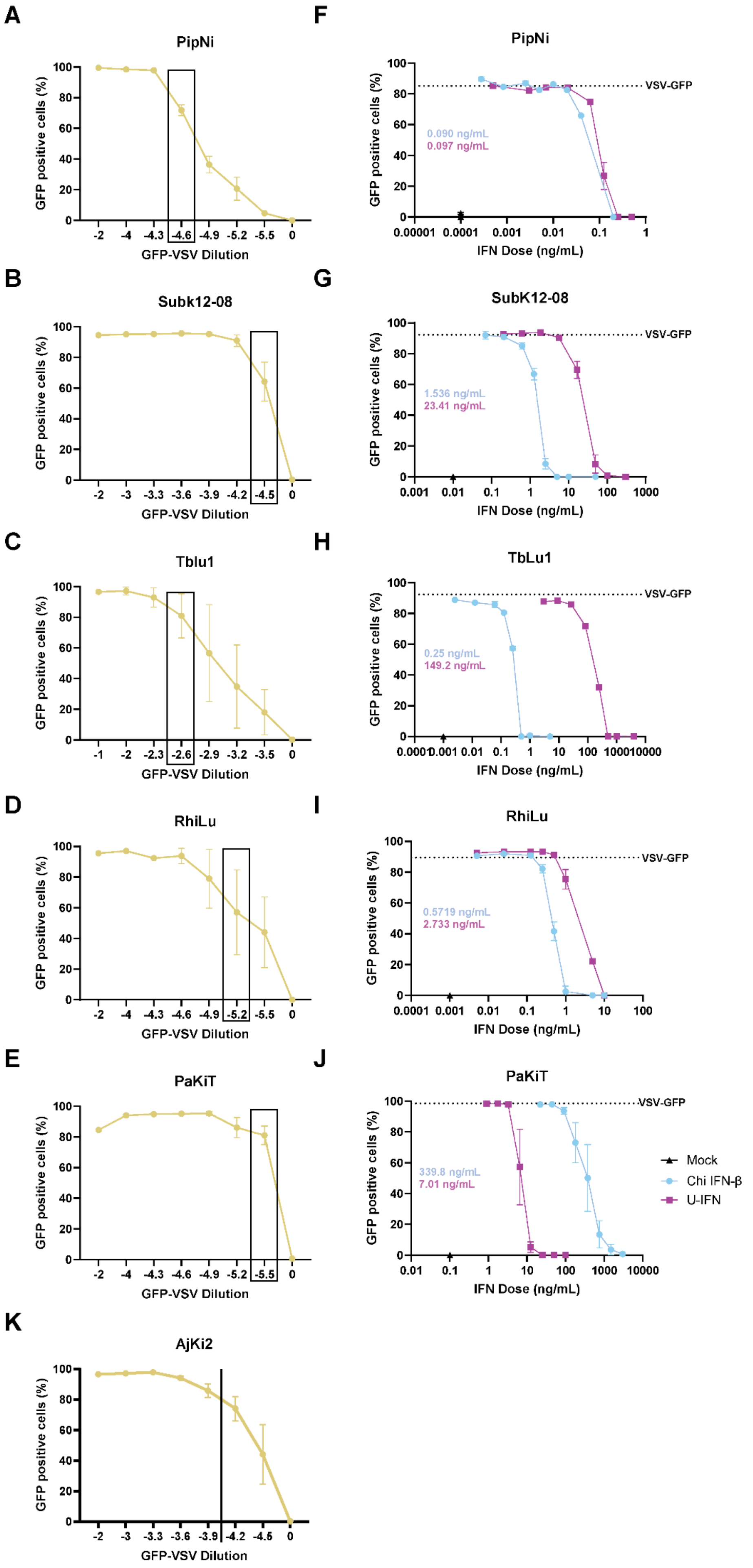
A
PipNi
GFP positive cells (%)
GFP-VSV Dilution
-2
-4
-4.3
-4.6
-4.9
-5.2
-5.5
0
B
Subk12-08
-2
-3
-3.3
-3.6
-3.9
-4.2
-4.5
0
C
Tblu1
-1
-2
-2.3
-2.6
-2.9
-3.2
-3.5
0
D
RhiLu
-2
-4
-4.3
-4.6
-4.9
-5.2
-5.5
0
E
PaKiT
K
AjKi2
-2
-3
-3.3
-3.6
-3.9
-4.2
-4.5
0
F
PipNi
VSV-GFP
0.090 ng/mL
0.097 ng/mL
IFN Dose (ng/mL)
G
SubK12-08
1.536 ng/mL
23.41 ng/mL
H
TbLu1
0.25 ng/mL
149.2 ng/mL
I
RhiLu
0.5719 ng/mL
2.733 ng/mL
J
PaKiT
339.8 ng/mL
7.01 ng/mL
Mock
Chi IFN-β
U-IFN

**Supplementary Fig 5. Chiroptera IFN-β induces an antiviral state in cell lines from diverse bat taxa**

(A-E) VSV-GFP doses yielding approximately 70–90 percent (%) GFP-positive single cells for each cell line were selected: PipNi ($10^{-4.6}$) (A), SubK12-08 ($10^{-4.5}$) (B), TbLu-1 ($10^{-2.6}$) (C), RhiLu ($10^{-5.2}$) (D) cells, and 1 PaKiT cells ($10^{4.6}$) (E). (F-J) Flow cytometry was used to quantify the percentage of single cells positive for GFP following treatment with a dilution series of IFNs and a single dose of VSV-GFP. IC50 values were calculated for Chi IFN-β and U-IFN for each cell line. Presented data a representative of at least three independent experiments conducted in quadruplicate. These data were used to calculate the fold difference in IC50 presented in Fig. 3O.

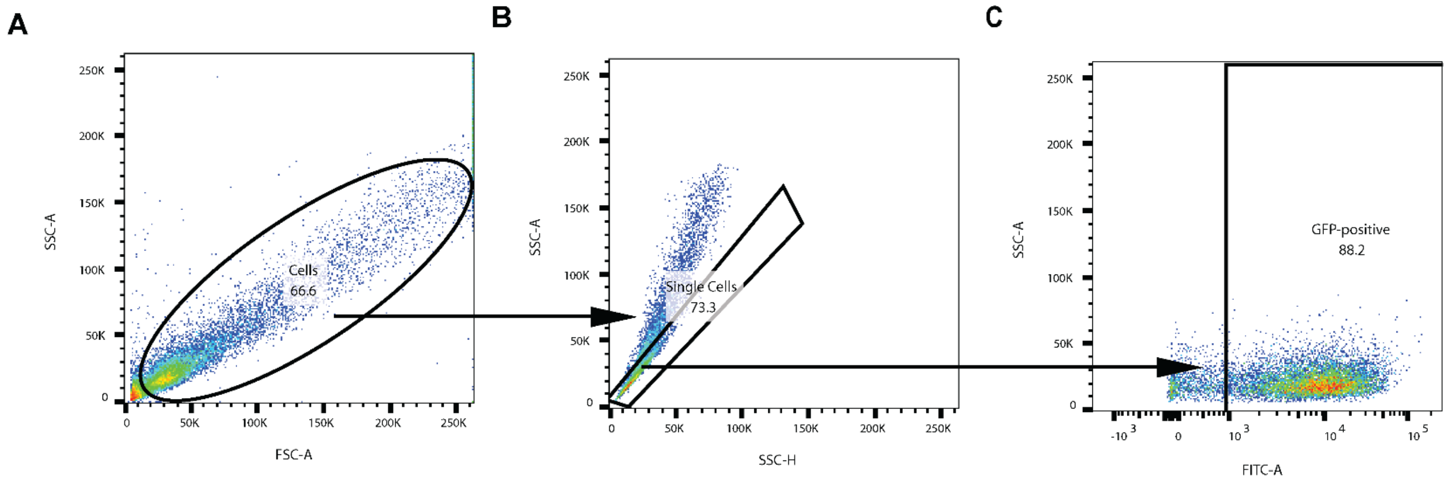


**Supplementary Fig 6. Gating strategy to evaluate infection with VSV- or EBOV-GFP**

Representative gating strategy for antiviral assays. (A) Cells were gated on SSC-A vs FSC-A to exclude debris and dead cells. (B) A SSC-A vs SSC-H gate was applied to limit analysis to single cells. (C) Singlets were then gated on FSC-A vs FITC-A to determine the percent of VSV-GFP-positive single cells. Strategy was applied to data presented in Fig 1G-H and Supp Fig 5A-J.
